# The intracellular region of truncated neurotrophin receptor TrkB-T1 promotes stroke-related effects in glial reactivity and neurotoxicity

**DOI:** 10.64898/2026.03.12.711279

**Authors:** Lola Ugalde-Triviño, María C. Serrano, Margarita Díaz-Guerra

**Affiliations:** Instituto de Investigaciones Biomédicas Sols-Morreale (IIBM), Consejo Superior de Investigaciones Científicas-Universidad Autónoma de Madrid, Madrid 28029, Spain; Instituto de Ciencia de Materiales de Madrid (ICMM), Consejo Superior de Investigaciones Científicas, Madrid 28049, Spain

**Keywords:** cell-penetrating peptides, excitotoxicity, gliosis, intranasal administration, neurodegeneration, neuroinflammation, NMDARs, stroke, TrkB-T1

## Abstract

The development of advanced therapies for stroke, spinal cord injury or neurodegenerative diseases –main causes of death, disability and dementia– requires a profound understanding of the complex interactions established among excitotoxic neuronal death, aberrant neurotrophic-signaling, glial reactivity, and neuroinflammation. However, the master proteins coordinating these mechanisms have not been yet defined. Different evidence suggests that the truncated form of the neurotrophin tyrosine kinase receptor, TrkB-T1, might play such a key role. The levels of this TrkB isoform increase in stroke while those of the full-length pro-survival isoform (TrkB-FL) are reduced. Additionally, ischemic stroke and, specifically, excitotoxicity induce TrkB-T1 regulated intramembrane proteolysis (RIP), a process releasing a receptor ectodomain able to bind the brain-derived neurotrophic factor (BDNF) and leading to decreased BDNF-signaling. We hypothesize that the second RIP product, TrkB-T1 intracellular domain (TrkB-T1-ICD), might similarly contribute to neurotoxicity but also reactive gliosis and neuroinflammation. Herein, we first demonstrate migration of the cytoplasmic TrkB-T1-ICD to the nuclei of neurons undergoing excitotoxicity, suggesting a possible role in the transcriptional control induced by injury. Then, taking advantage of cell-penetrating peptides (CPPs), we produce a TrkB-T1-ICD mock peptide (Bio-LTT1_Ct_) containing the short TrkB-T1 intracellular region (23 amino acids) and test it *in vitro* and *in vivo*. Notably, this peptide migrates to the nucleus of both neurons and astrocytes cultured *in vitro* and provokes cell death. Additionally, Bio-LTT1_Ct_ induces early transcriptional changes in neurons resembling those triggered by excitotoxicity such as the inhibition of the promoter activity of pro-survival transcription factors CREB and MEF2, and altered mRNA levels of their regulated genes. *In vivo*, Bio-LTT1_Ct_ is accessible to the brain cortex after intranasal delivery, being efficiently distributed into cortical neurons and astrocytes of both hemispheres. Moreover, peptide administration is sufficient to promote important pathological hallmarks of stroke such as the imbalance of the TrkB isoforms, and the reactivity of astrocyte and microglia, cells that acquire proinflammatory profiles. Altogether, these results establish TrkB-T1 RIP as a central mechanism of ischemic damage and demonstrate that the receptor intracellular region is sufficient to recapitulate stroke-like effects on neurotoxicity, glial reactivity and neuroinflammation.

## Background

Stroke is the second leading cause of death worldwide (≈ 7 million deaths per year, representing about an 11% of the total) and an important cause of dementia and acquired disability among the 94 million of survivors (1). Specifically, ischemic stroke (≈ 85% of the total cases) results from the occlusion of a cerebral blood vessel which leads to severe focal brain hypoperfusion. This insult is followed by nutrient and oxygen deprivation, causing neuronal death and irreversible damage to the affected tissue, known as the infarct core. Surrounding this area is the penumbra, which experiences a less severe decrease of cerebral blood flow and, although functionally impaired, remains metabolically active. Since this tissue can persist viable for several hours, it has become an important target for neuroprotection. However, if the blood flow is not restored or no therapy is applied, neurons in this area can suffer secondary death causing the expansion of the infarct core. This delayed death is mainly due to excitotoxicity, promoted by glutamate accumulation in the extracellular space and the consequent overactivation of its receptors, particularly N-methyl-D-aspartate receptors (NMDARs), which is followed by massive Ca^2+^ influx and ion homeostasis disruption. Additional pathological processes in ischemic stroke are blood-brain barrier (BBB) disruption and neuroinflammation, which exacerbates brain injury in the acute phase of stroke (2). However, inflammation also exerts beneficial effects during the recovery phase by facilitating brain repair processes (3, 4). Microglial activation, which occurs within minutes and results in transcriptional regulation and prominent morphological changes, engages in crosstalk with astrocyte activation through cytokine secretion (5). Reactive astrogliosis, a process that takes place in the peri-infarct environment, also implies a dramatic morphological transformation, enhanced proliferation/migration and upregulation of genes such as that encoding for the glial fibrillary acidic protein (GFAP) (6). Specifically, some reactive astrocytes lose their normal functions, including promotion of neuronal survival and outgrowth, and gain a pro-inflammatory phenotype that triggers detrimental effects on neuronal and oligodendrocyte survival (7–9). Within this context, recent data suggest that the truncated tropomyosin-related kinase B receptor (TrkB-T1), the most abundant CNS TrkB isoform in adults, might play a key role in the crosstalk between excitotoxicity, reactive gliosis and neuroinflammation in stroke.

Under physiological conditions, alternative splicing of the *Ntrk2* gene produces different TrkB isoforms (10), full-length TrkB (TrkB-FL) and truncated TrkB-T1 being the major murine cortical isoforms (11). While TrkB-FL is mostly expressed in neurons (12), TrkB-T1 is mainly produced by astrocytes and, to a lesser extent, neurons. Binding of brain-derived neurotrophic factor (BDNF) to TrkB-FL induces receptor dimerization, tyrosine kinase (TK) activity, transphosphorylation, and activation of interrelated signaling pathways that, altogether, promote pivotal CNS processes such as neuronal survival and synaptic function. An important downstream mechanism of BDNF/TrkB-FL-signaling is the activation of pro-survival transcription factors (TFs), cAMP response element-binding protein (CREB) (13) and myocyte enhancer factor 2 (MEF2) (14, 15). They regulate many different genes, including those encoding BDNF (16–18), TrkB (19) and some NMDAR subunits (20). In neurons, TrkB-T1, which lacks the TK domain but can still bind BDNF, acts as a TrkB-FL dominant-negative receptor inhibiting neurotrophic-signaling (21). However, TrkB-T1 also exerts TrkB-FL-independent actions, including fine-tuning of neurotrophic-signaling (22) and regulation of astrocyte functions such as calcium release from intracellular stores (23), neurotransmitter transport (24, 25), and cell maturation and morphology (26, 27). A long-standing idea is that TrkB-T1 individual actions rely on protein interactions established by its highly conserved and isoform-specific C-ter sequence (FVLFHKIPLDG) (11, 21). In fact, we have recently isolated and characterized the TrkB-T1 isoform-specific interactome, proving interactions with proteins related to gene expression and mRNA regulation, including splicing and translation (28).

In stroke, neurotrophic support becomes profoundly aberrant, partly due to the unbalance of the TrkB isoforms induced by excitotoxicity (29–31). TrkB-FL levels decrease in the infarct core while TrkB-T1 increases in surrounding astrocytes (32), promoting brain injury and contributing to edema formation (33). These changes start very early after ischemia, the TrkB-T1 upregulation being concurrent with increased GFAP expression in astrocytes of the emerging fibroglial scar. Moreover, most GFAP^+^ cells also express complement component 3 (C3), a marker of inflammatory reactive astrocytes. Later on, the number of GFAP^+^/C3^+^ cells highly increases, together with macrophage infiltration and microglial activation at the infarct border and adjacent tissue (28). Similar TrkB-T1 upregulation takes place in neurodegenerative diseases (NDDs) (31), Down syndrome (34) and spinal cord injury (SCI) (35). Notably, the restoration of TrkB-T1 physiological levels rescues neuronal cell death in a Down syndrome model (34), while a TrkB-T1 specific deletion in astrocytes facilitates functional recovery in SCI (36). In this context, we have recently demonstrated the importance of the TrkB-T1 isoform-specific interactions in the promotion of stroke-associated reactive astrogliosis and neuroinflammation. Specifically, we have designed a TrkB-T1-derived cell-penetrating peptide (CPP), denominated TT1_Ct_, formed by two elements: a short basic sequence derived from the HIV Tat protein, having the ability to cross the plasma membrane and the BBB, and the TrkB-T1 isoform-specific C-ter sequence. Peptide TT1_Ct_ not only interferes with reactive gliosis and neuroinflammation after ischemia, but also reduces infarct size and neurological damage (28), suggesting that TrkB-T1 might indeed act as a master regulator of these interconnected pathological processes in stroke, as well as in other acute and chronic CNS diseases similarly associated with excitotoxicity (37).

Additional mechanisms contribute to aberrant BDNF/TrkB signaling in excitotoxicity and ischemia: (a) retrograde TrkB-FL transport to the Golgi apparatus where it is processed by calpain, producing a truncated receptor similar to TrkB-T1 (29, 38–40); (b) changes in the profile of TrkB-T1 isoform-specific interactions, including upregulation of pathways important for the promotion of neurotoxicity and astrocyte reactivity (28); and (c) activation of regulated intramembrane proteolysis (RIP), which involves the sequential processing of both TrkB isoforms by metalloproteases, shedding identical extracellular domains (ECDs), followed by γ-secretase proteolysis of the remaining transmembrane fragments and the release of the intracellular domain (ICD) into the cytoplasm (38). Excitotoxicity-induced RIP is a major mechanism of TrkB-T1 regulation but only secondarily controls TrkB-FL. In general, this type of proteolysis contributes to neuronal function under both physiological and pathological conditions. For instance, RIP is important for the regulation of synaptic function (41), while the dysregulation of metalloproteases or γ-secretase activities is involved in relevant neurological disorders (42, 43). In previous work, we proposed that the TrkB-T1 RIP products generated in excitotoxic conditions might notably contribute to ischemic injury. Actually, we have already demonstrated that TrkB-ECD is capable of BDNF binding and acts as a BDNF-scavenger, thus having a deleterious effect on BDNF/TrkB signaling (38). However, similarly to most γ-secretase substrates, the role played by TrkB-T1-ICD in ischemia has not yet been characterized. Once produced, the soluble ICDs might be further degraded or act as signaling molecules, playing local functions in the cytosol near their cleavage site or, more often, being translocated to the cell nucleus where they modulate gene expression. For example, Notch-ICD partly mediates Notch receptor effects on learning and memory by repressing CREB-dependent gene transcription (44). In this work, we have first characterized the formation and fate of TrkB-T1-ICD after NMDAR overactivation in primary cultures of cortical neurons and astrocytes. Then, by taking advantage of CPP properties, we have produced a TrkB-T1-ICD mock peptide which has been tested both *in vitro* and *in vivo*. The results obtained demonstrate that the short TrkB-T1 intracellular region is sufficient to induce by itself the imbalance of the TrkB isoforms and the activation of glial cells, which acquire a proinflammatory profile, resembling important pathological stroke-related mechanisms. Additionally, our data strongly suggest that TrkB-T1 RIP is a main mechanism of brain injury after stroke, affecting both neurons and glial cells.

## Materials and Methods

Reagents and resources are described as supplementary information, additional file 1.

### Synthetic peptides

Synthetic cell-penetrating peptides (> 95% purity; GenScript) were used for treatment of primary cortical cultures (25 µM; *in vitro* studies) and mice (2.5 nmol/g; *in vivo* studies). All these CPPs contained a HIV-1 Tat sequence (11 amino acids) linked to a specific TrkB-T1 or c-Myc sequence as indicated (see supplementary information, additional file 1). Additionally, these peptides contained a biotin molecule at the N-ter, which helped peptide visualization and stabilization, and an amine group at their C-ter. Peptides were directly added to the culture media for *in vitro* experiments or intranasally administered in *in vivo* tests.

### Experimental models

Animal procedures were performed following European Union Directive 2010/63/ EU and were approved by the CSIC and Comunidad de Madrid (Ref PROEX 276.6/20) ethics committees. The housing facility was approved by Comunidad de Madrid (#ES 280790000188) and conforms to official regulations. Animals had standard health and immune status and were looked after daily by professional caretakers. Male mice were kept in groups of up to five animals in standard IVC cages while 1–2 pregnant rats occupied standard cages, always containing bedding and nesting materials. Animals were under controlled lighting (12 h light cycles), relative humidity and temperature conditions. Irradiated food and water were provided *ad libitum*. The three Rs principle was carefully respected.

### Primary cultures of rat cortical neurons and induction of excitotoxicity

Dissected cerebral cortices of Wistar rat embryos (E18; both genders included) were mechanically dissociated in culture medium (Minimum Essential Medium, MEM, Life Technologies) followed by seeding of the cell suspension at a density of 1.1×10^6^ cells/ml in MEM supplemented with 22.2 mM glucose, 0.1 mM glutamax, 5% fetal bovine serum (FBS), 5% horse serum (HS), 100 U/ml penicillin, and 100 µg/ml streptomycin. Before seeding, plates were treated overnight with 100 µg/ml poly-L-lysine and 4 µg/ml laminin at 37°C. Cells were incubated in an atmosphere of 5% CO_2_ and 95% humidity at 37°C. In these cultures, glial growth was repressed after 7 days *in vitro* (DIVs) by adding 10 µM cytosine β-D-arabinofuranoside (AraC) and experimental treatments added after 12 DIVs. These cultures presented a high percentage of mature neurons, combined with a small proportion of glial cells, mostly astrocytes (29), which help to sustain non-toxic glutamate concentrations in the medium. A robust *in vitro* excitotoxic response was induced in these cultures by incubation with high concentrations of the glutamate analog NMDA (100 µM) and co-agonist glycine (10 µM), herein denoted simply as NMDA. This treatment causes neuronal death but has no effect on astrocyte viability (29). When indicated, cultures were preincubated with TAPI-2 (10 µM) for 30 min before NMDA addition to the media.

### Primary cultures of rat cortical astrocytes and treatment

Dissected cerebral cortices obtained from Wistar rat embryos (E18) as described before were mechanically dissociated in Hank’s Balanced Salt Solution (HBSS, Thermo Fisher) containing CaCl₂ and MgCl₂, supplemented with 10 mM HEPES. The resulting cell suspension was then seeded in 75 cm² flasks pretreated with 100 μg/mL poly-L-lysine at a density of 4×10⁶ cells/ml in proliferation medium. This medium consisted of Dulbecco’s Modified Eagle Medium/Nutrient Mixture F-12 with GlutaMAX (DMEM/F-12/GlutaMAX, Thermo Fisher), 10% FBS, 5% HS, 100 U/ml penicillin, 100 μg/ml streptomycin, 2% B27 supplement, 10 ng/ml epidermal growth factor (EGF), and 10 ng/ml basic fibroblast growth factor (bFGF). After 4 DIVs, when cultures reached approximately 70–80% confluence, the medium was removed, and the flasks were vigorously washed with PBS to eliminate contaminating oligodendrocytes. Cell cultures were then trypsinized using TrypLE Express (Thermo Fisher) and reseeded in flasks under the same culture conditions at a density of 3×10⁶ cells/ml in proliferation medium. Six days later, astrocytes were washed, trypsinized as before, and seeded at a density of 0.7×10⁶ cells/ml in poly-L-lysine-coated plates for 24 h. After this period, the plates were washed with PBS, and the medium was replaced with differentiation medium consisting of DMEM/F-12/GlutaMAX, 10% FBS, 100 U/ml penicillin, and 100 μg/ml streptomycin. Twenty-four hours later, cultures were treated with 400 μM NMDA or 100 μM APMA for selected times.

### Mice model of brain damage by Bio-LTT1_Ct_ administration

Adult male mice were treated with a single dose (2.5 nmol/g) of Bio-TMyc or Bio-LTT1_Ct_ peptides, administered intranasally into one nostril to mildly sedated animals. After administration, animals were sacrificed 1 or 48 h later and intracardially perfused for immunohistochemistry analysis.

### Assessment of neuronal injury in cortical cultures

Thiazolyl blue formazan (MTT) reduction assay was used to determine cell viability. At corresponding time points, 0.5 mg/ml MTT was added to the medium and incubated for 2 h at 37°C. The formazan salts formed were then solubilized using DMSO. The results were quantified spectrophotometrically at 570 nm. This procedure was employed to establish the effects on total cell viability of treatments with Bio-LTT1_Ct_ (cultures of either neurons or astrocytes) or NMDA (cultures of astrocytes). Only for the analysis of RIP contribution to excitotoxicity, we specifically calculated neuronal viability. As mentioned before, neuronal cultures contained a small percentage of glial cells, resistant to the excitotoxic damage (45, 46). Therefore, we needed to calculate the contribution of glial viability to the total values. Thus, 24 h before the MTT assay, sister cultures were treated with 400 µM NMDA and 10 µM glycine, conditions that induced a complete neuronal death but no glial damage. The obtained absorbance value (glial viability) was then subtracted from total values to calculate the specific viability of neurons present in the cultures.

### Subcellular fractionation

Cell cultures were washed twice at 4°C with Tris Natrium-EDTA buffer (TNE, 150 mM NaCl, 50 mM Tris HCl pH 7.5, and 1 mM EDTA) and collected in the same buffer. A small aliquot of cell suspension was separated to prepare the total lysate (TL), while the remaining was used for cell fractionation to obtain the post-nuclear (Post-N) and nuclear (N) phases. The TL was prepared in NPBD buffer (10 mM NaCl, 10 mM Tris HCl pH 7.5, 2 mM MgCl_2_, 0.5% NP40, 0.1% SDS, and 1% sodium deoxycholate), containing protease and phosphatase inhibitors as before. After a 30 min incubation at 4°C, the lysate was sonicated using a Bioruptor apparatus (Diagenode) and the cellular debris were eliminated by centrifugation at 3000 rpm for 10 min at 4°C to obtain the final protein suspension. The Post-N phase was obtained by mild cell lysis with NPB buffer (10 mM NaCl, 10 mM Tris HCl pH 7.5, 2 mM MgCl_2_, and 0.5% NP-40), containing 2 mM DTT and protease and phosphatase inhibitors, conditions that maintained the nuclear integrity. After a 30 min incubation at 4°C, the nuclei were separated from the Post-N fraction by centrifugation. Once the supernatant was collected (Post-N), the nuclei were washed twice with NPB buffer with 2 mM DTT and the inhibitors, and then lysed with NPBD buffer, followed by sonication. After removal of nuclear debris by centrifugation, the N phase was collected. All fractions were analyzed by BCA to determine protein concentration, fractioned by SDS-PAGE and analyzed by immunoblot.

### Immunoblot analysis

Cells were lysed in RIPA buffer (50 mM Tris-HCl pH 8.0, 150 mM NaCl, 1% sodium deoxycholate, 1% NP-40, 0.1% SDS, and 1 mM DTT), supplemented with protease and phosphatase inhibitors (see supplementary information, additional file 1), for 30 min at 4°C. Then, cell lysates were sonicated using a Bioruptor apparatus and centrifugated at 4°C for 20 min at 11,000 rpm. The protein concentration in lysates was measured by using a BCA Protein Assay Kit (Thermo Fisher) followed by 5 min denaturalization in SDS-sample buffer at 95°C. Equal amounts of total lysates were resolved in Tris-Glycine SDS-PAGE and transferred to a nitrocellulose membrane (GE Healthcare) in 25 mM Tris HCl pH 8.3, 250 mM glycine and 10% methanol, using an electric current of 400 mA for 70 min. After transfer, membranes were stained with a Ponceau S solution to check efficacy. Then, membranes were blocked for 30 min with a 5% non-fat dry milk solution in Tris Buffered Saline-Tween (TBS-T, 20 mM Tris HCl pH 7.5, 137 mM NaCl, and 0.05% Tween-20) and incubated overnight at 4°C with primary antibodies (see supplementary information, additional file 1). Next, membranes were washed with TBS-T and incubated with the appropriate anti-rabbit or anti-mouse peroxidase-conjugated secondary antibodies (Bethyl) for 1 h. Immunoreactivity was detected using Clarity Western ECL Blotting Substrate (BioRad) and band intensity was quantified by densitometric analysis (Photoshop, Adobe). Levels of GFAP were normalized using those of β-actin present in the same sample and expressed relative to the maximum values obtained in each experiment, arbitrarily given a 100% value.

### Cell transfection and gene reporter assays

Plasmids used contained minimal CREB or MEF2 response elements upstream firefly luciferase reporter genes (respectively, pCRE or pMEF2; see details in supplementary information, additional file 1). Primary cultures, without AraC treatment, were transfected at 11 DIVs with 0.4 µg of those plasmids using Lipofectamine 2000. DNA-liposomes complexes were prepared according to the manufacturer’s instructions in neurobasal medium (Thermo Fisher Scientific) with 1 mM glutamax (Gibco, Thermo Fisher Scientific) and added to the cell cultures. After 2 h of transfection, this mix was replaced with previously collected conditioned medium and incubation proceeded to complete 24 h. Protein extracts were obtained using Passive Lysis Buffer (Promega, Cat# E1941) and luciferase activity was detected by using a luminometer (GloMax 96 microplate luminometer, Promega), samples being in REAP buffer (25 mM glycylglycine, 15 mM SO_4_Mg, 4 mM EGTA, 15 mM potassium phosphate pH 7.8, 3.3 mM ATP, 1 mM DTT, and 75 µM luciferin).

### RNA extraction and qPCR assay

Total RNA was extracted using QIAcube technology and treated with DNases before cDNA synthesis by the High Capacity cDNA Reverse Transcription Kit (Applied Biosystems). A 7900HT Fast real-time PCR system (Applied Biosystems) was used for SYBR green gene expression assays (primers indicated in the supplementary information, additional file 1). PCR conditions were 10 min at 95°C, followed by 40 cycles of 15 s at 95°C and 60 s at 60°C. For each independent experiment, we made a specific standard curve for every gene and technical triplicates were prepared for every sample. Results were normalized to the levels of NSE or GAPDH genes as indicated.

### Peptide visualization in primary neural cultures and immunocytochemistry

Primary cultures, seeded at half of the concentration previously indicated, were grown for 13 DIVs on coverslips previously treated overnight at 37°C with poly-L-lysine and laminin as described before. In experiments designed for peptide visualization, cells were incubated with Bio-TMyc or Bio-LTT1_Ct_ (25 µM, 30 min), or left untreated, and then fixed for 30 min with 4% paraformaldehyde in PBS. The fixed cells were washed several times with PBS before they were blocked and permeabilized for 4 h with 4% goat serum and 0.5% Triton X-100 prepared in PBS. Afterwards, coverslips were incubated overnight at 4°C with the TrkB-T1 isoform-specific antibody diluted in the same blocking solution. After washing with PBS, the coverslips were incubated with the corresponding secondary antibody together with Fluorescein Avidin D (200 µg/ml) for 1 h at room temperature and then 10 min with DAPI to stain DNA. Finally, the samples were mounted in Prolong Diamond. Images were acquired using an inverted Zeiss LSM 710 laser confocal microscope (Jena, Germany) with 40x and 63x Plan-Apochromatic oil immersion objectives. Images correspond to single sections or maximum intensity projections and were normalized for each color separately and processed with ImageJ (NIH Image) and Fiji. For immunocytochemistry of cultures treated with NMDA, cells were fixed as described before and incubated with anti-TrkB-T1 antibodies alone (astrocytes) or together with anti-NeuN antibodies (neurons). Then, the procedure continued as indicated above but without incubation with Fluorescein Avidin D.

### Peptide visualization in mice brain and immunohistochemistry

After intranasal (*i.n.*) peptide administration for 1 or 48 h, animals were deeply anesthetized and intracardially perfused with cold PBS and 4% paraformaldehyde in PBS. Brains were postfixed in the same fixative at 4°C for 24 h and cryoprotected in 30% sucrose for 48 h at 4°C. Floating coronal frozen sections (30 µm thick) obtained using a cryostat (Leica, Heidelberg, Germany) were incubated in blocking solution (10% goat serum, 5% BSA, and 0.5% Triton X-100 in PBS) for 3 h at room temperature. When indicated, sections were incubated overnight with NeuN in 4% goat serum right after blocking and, after washing, with Alexa Fluor 488 or 647-conjugated antibodies together with Cy5-Streptavidin (Cy5-Strep, 5 µg/ml) or Fluorescein Avidin D (FluorAv, 200 µg/ml) and DAPI (5 µg/ml) for 1 h. Finally, sections were mounted and dried on slides, and cover slipped with Prolong Diamond. Image acquisition was performed using a Cell Observer (Zeiss), for wide-field peptide imaging in whole coronal sections labeled with Cy5-Streptavidin, or with an inverted laser confocal microscope with a 63x Plan-Apochromatic oil immersion objective for those labeled with FluorAv. Images presented were acquired and processed equally for each condition and corresponded to single sections. Background was subtracted using vehicle-injected animals.

For immunohistochemistry, brains were processed and cryoprotected as described before. Coronal frozen sections were incubated in flotation with the blocking solution for 1 h at room temperature, followed by an overnight incubation with the indicated primary antibodies diluted in 4% goat serum and 0.3% Triton X-100 in PBS at 4°C. After washing, slides were incubated 1 h with secondary antibodies (see supplementary information, additional file 1) and DAPI (5 μg/ml). Sections were then washed in PBS, mounted on slides, dried for 15 min at 37°C on a hot plate or air dried overnight, and then cover slipped with Prolong Diamond. Confocal images were acquired with 40x and 63x Plan-Apochromatic oil immersion objectives and processed as described above.

Images used for Iba1-based morphological analysis were obtained using an inverted Stellaris 8 Leica confocal microscope with a 40x objective. A tile of approximately 12 x 6 fields from 30 µm-thick brain coronal sections, corresponding to the cortical area, was acquired. The z-stacks covered the sample width, with 40% overlap between stacks. AIVIA software version 15.0.0.42239 was used for 3D reconstruction, segmentation and morphological measurements. The segmentation code used for the analysis is available as supplementary information (see additional file 5).

### Statistical analysis

Data are expressed as mean ± standard error of the mean (SEM) of at least three independent experiments in all cases (N ≥ 3). For viability, mRNA and gene reporter assays, technical replicates were included in each independent experiment. Specific details of the N and n values, and the statistical test applied in each case can be found in the respective legends. The assignment of treatment was performed randomly. Statistical analyses were performed in GraphPad Prism 8.0.2. and RStudio 2024.04.2. The normality of the data was analyzed by Saphiro-Wilk test. In all cases, significance was defined as: \**P* < 0.05, ** *P* < 0.01, *** *P* < 0.001, and **** *P* < 0.0001. A *P*-value larger than 0.05 was considered as non-significant (n.s.).

## Results

### Characterization of TrkB-T1-ICD formation and intracellular localization in primary cultures of cortical neurons and astrocytes

The precise TrkB-T1 intramembrane sequence cleaved by γ-secretase has not yet been defined (Fig. 1A, left panel). We hypothesize that the resulting TrkB-T1-ICD would be formed by two elements: an undetermined stretch of the TrkB-T1 transmembrane (TM) sequence and the short intracellular C-ter region (23 amino acids), composed of a sequence shared with TrkB-FL (amino acids 454-465) and a TrkB-T1-isoform specific sequence (amino acids 466-476; Fig. 1A, right panel). For these studies, we used a TrkB-T1-specific antibody, prepared using the isoform specific sequence as an antigen and able to detect the TrkB-T1-ICD, in addition to the complete isoform (95 kDa). Using this antibody, we first analyzed the kinetics of fragment formation and its subcellular localization in primary cultures of cortical neurons subjected to excitotoxicity (Fig. 1B, C). Overactivation of NMDARs with their co-agonists NMDA and glycine (combined treatment hereafter abbreviated as NMDA) induced the formation of a low molecular weight receptor fragment in total lysates (TL), in parallel to a decrease of total CREB levels (tCREB) previously described to be associated with excitotoxic conditions (39). By cell fractionation, we demonstrated that fragment TrkB-T1-ICD migrated to the cell nucleus where it progressively accumulated, differences being statistically significant after 6 h of NMDA treatment (\*\**P* < 0.01 with respect to basal conditions, Fig. 1C). Immunofluorescence experiments showed a basal TrkB-T1 presence in neurons, counterstained with the specific marker NeuN (arrowheads) and main cell subpopulation in these cultures, but also in NeuN^-^cells (asterisk), corresponding to a minor astrocyte subpopulation (Fig. 1D). Indeed, the levels of TrkB-T1 receptor were much higher in astrocytes compared to neurons, as previously described (28). A short treatment with NMDA (40 min) was able to induce partial migration of a polypeptide recognized by the TrkB-T1 antibody, presumably TrkB-T1-ICD, to the nucleus of NeuN^+^ neurons.

**Fig. 1.**
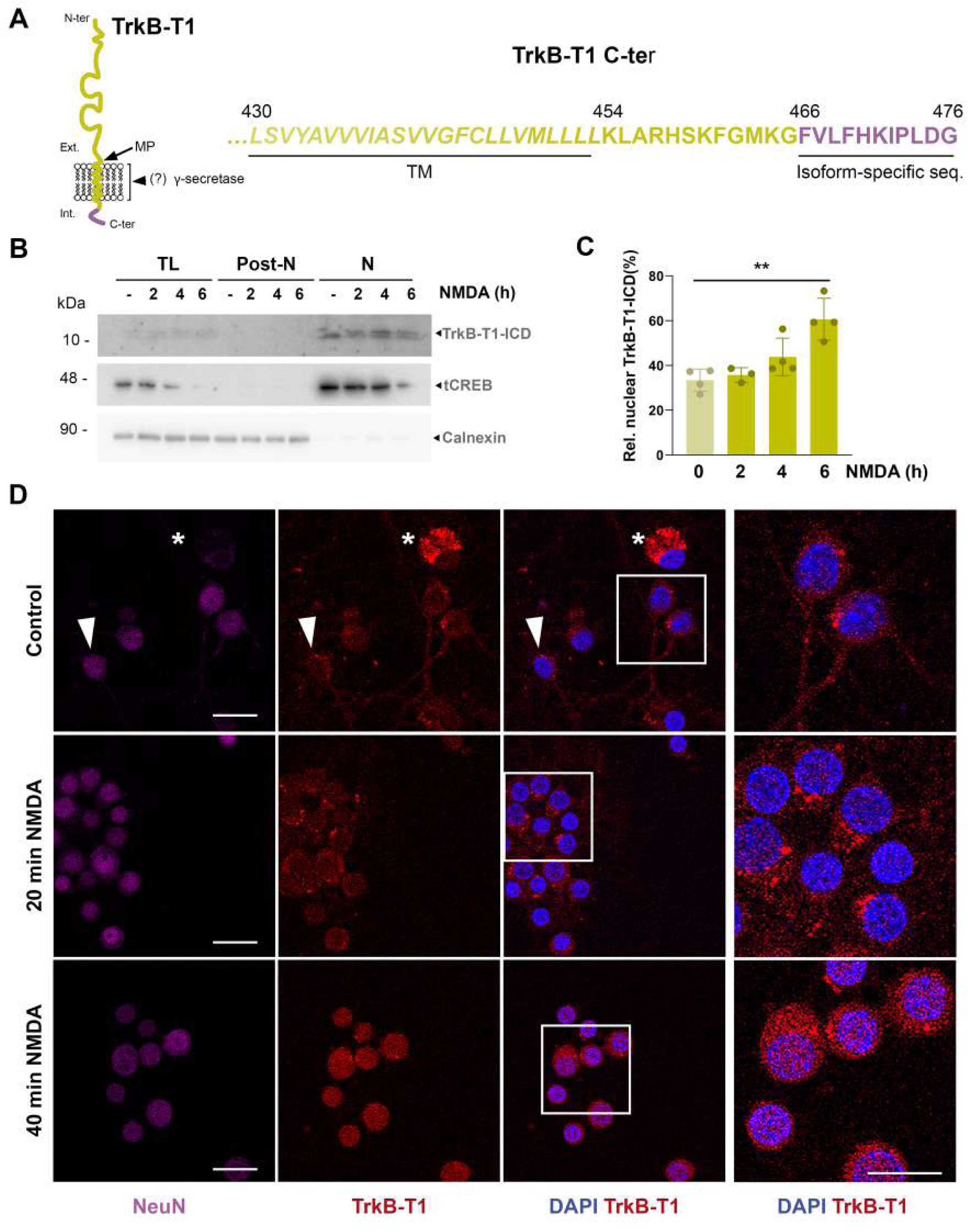
TrkB-T1 RIP induced by excitotoxicity leads to TrkB-T1-ICD formation and nuclear migration in neurons. **A** Scheme of TrkB-T1 membrane orientation with sites of metalloproteinase (MP) and γ-secretase cleavage, located in undefined extracellular and intramembrane amino acids, respectively. The TrkB-T1 C-ter sequence is also shown, with indication of the transmembrane (TM) region (*italics*) and the intracellular sequence, which contains the isoform-specific sequence (pink). **B** Subcellular fractionation of primary cultures of neurons and astrocytes treated for 2, 4 or 6 h with NMDA (100 µM) and glycine (10 µM), herein denoted simply as NMDA. A low molecular weight C-ter fragment, probably the TrkB-T1-ICD, was detected in the nuclear fraction (N) by the TrkB-T1 isoform-specific antibody. Correct cell fractionation was confirmed by using calnexin as a Post-nuclear (Post-N) marker and tCREB as a nuclear (N) marker. **C** Quantification by densitometric analysis of TrkB-T1-ICD nuclear levels. Levels of nuclear TrkB-T1-ICD for each time point were expressed relative to those corresponding to the maximum values obtained for that particular experiment (100%). Individual data and mean ± SEM (N = 4) are presented. Results were analyzed by one-way ANOVA test followed by *post hoc* Dunnett’s test. The comparison between the control and 6 h of NMDA treatment is statistically significant (\*\**P* = 0.0011). **D** Immunocytochemistry assay of cortical cultures in basal conditions and treated with NMDA for 20 or 40 min. TrkB-T1 distribution was visualized by labelling with the isoform specific antibody (red). Neurons (arrowheads) were identified using the specific marker NeuN (magenta) and nuclei were stained with DAPI (blue). Neu^-^/TrkB-T1^+^ cells (asterisk) were rare and probably correspond to a small astrocyte subpopulation present in these cultures. Representative confocal microscopy images are presented. Scale bar represents 20 µm in all images.

NMDARs are also expressed by glial cells, particularly astrocytes, although their function is poorly understood (47). Therefore, it was important to investigate if overactivation of astrocytic NMDARs also induced TrkB-T1 RIP. With that aim, we prepared primary cultures of cortical astrocytes, where TrkB-T1 levels are 7 times higher than those found in mixed cultures (28). We confirmed that high NMDA concentrations in these astrocyte-enriched cultures did not affect their viability even after long treatments (24 h, Fig. 2A), as previously described (45, 46). In contrast, this glutamate analog triggered intracellular cascades that promoted increased levels of TrkB-T1 (Fig. 2B) and GFAP (Fig. 2C, D). Nonetheless, we could not demonstrate the formation of TrkB-T1 RIP fragments, TrkB-ECD and TrkB-T1-ICD, upon NMDA treatment in these cultures. Only the direct activation of matrix metalloproteinases by p-aminophenylmercuric acetate (APMA) was able to induce TrkB-T1-ICD at very low levels, this fragment again concentrating in the cell nucleus (Fig. S1). Based on these results, we conclude that glutamate-induced TrkB-T1 RIP is restricted in astrocytes *in vitro* while being strongly induced in neurons exposed to excitotoxicity. Finally, in order to establish a possible TrkB-T1-ICD contribution to neuronal death, occurring jointly with that derived from TrkB-ECD shedding (38), we investigated if generic RIP inhibition was neuroprotective. Cultured neurons were pretreated with a broad-spectrum metalloprotease inhibitor (TAPI-2), blocking the first and necessary RIP step, before NMDA treatment for 1 or 2 h (Fig. S2). We found that RIP inhibition had a statistically significant effect on neuronal survival under excitotoxic conditions, as demonstrated by an increased neuronal viability in cultures treated with TAPI-2 + NMDA compared to those incubated only with NMDA (77 ± 2% and 54 ± 12% for 2 h of treatment, respectively, \**P* < 0.05). However, this preliminary result could not exclude potential neurotoxic events due to alternative metalloproteinase substrates different from TrkB-T1.

**Fig. 2.**
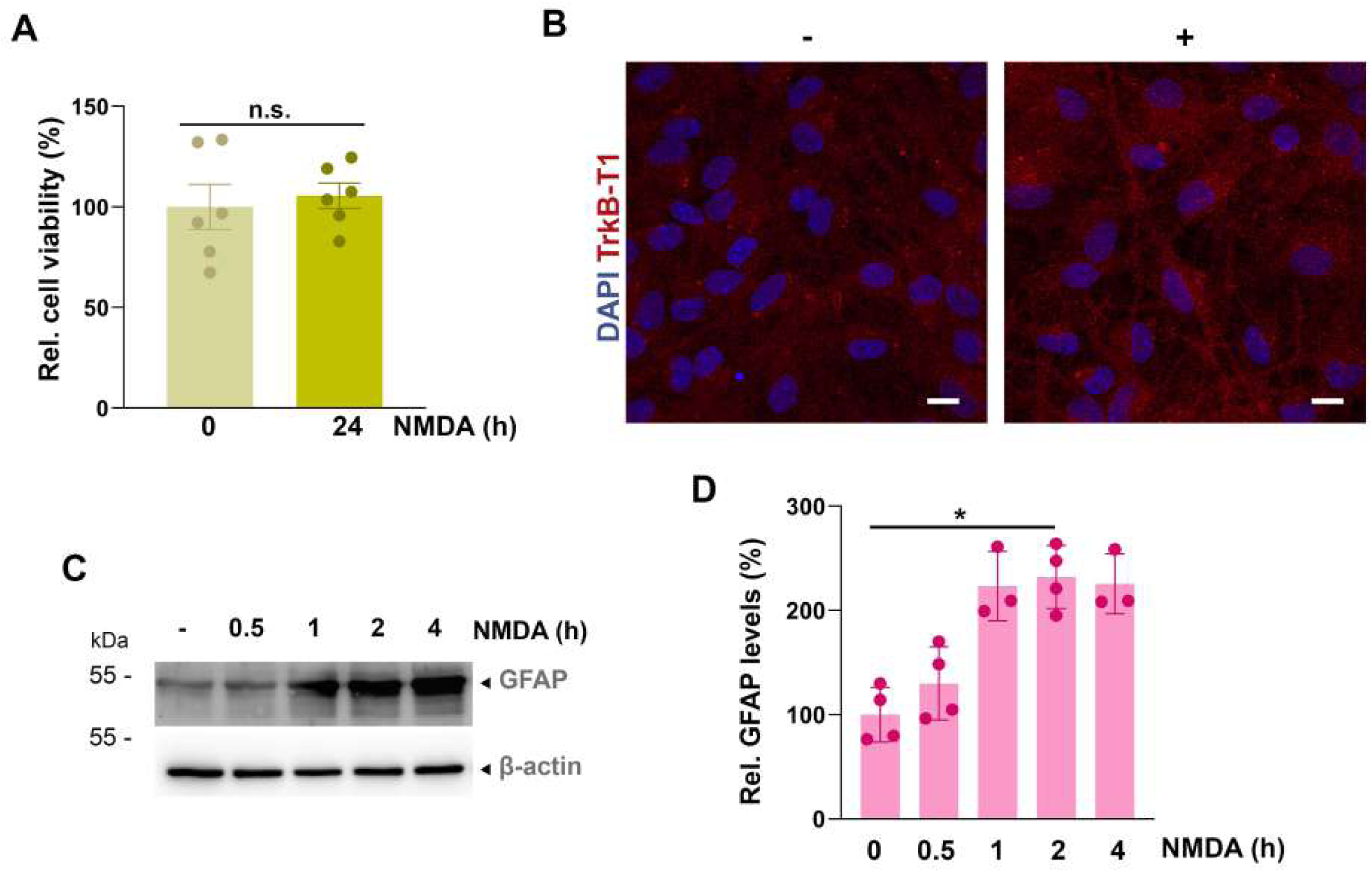
TrkB-T1 RIP is not activated in primary cultures of cortical astrocytes upon NMDA treatment. **A** Cell viability of cultured astrocytes treated with NMDA (400 µM) and glycine (10 µM) for 24 h. Individual data and mean ± SEM are presented relative to values obtained for untreated cells (N = 6). Technical replicates were included in each independent experiment (n = 3). Viability levels were analyzed by unpaired Student’s *t*-test, being not significantly different (n.s.). **B** Immunocytochemistry assay of cultured astrocytes treated with NMDA for 4 h (+ condition). Basal conditions are also shown for comparison. TrkB-T1 was detected by immunolabeling with an isoform-specific antibody (red) and cell nuclei using DAPI (blue). Representative confocal microscopy images are shown. Scale bar represents 10 µm. **C** Western Blot analysis of GFAP levels in astrocytes treated with NMDA as indicated for 0.5-4 h. β-actin levels were measured as loading control. **D** Quantification by densitometric analysis of GFAP levels. After normalization to β-actin expression, GFAP levels were expressed relative to those corresponding to basal conditions (assigned value of 100). Individual values are presented and data are expressed as the mean ± SEM (N = 3-4).

### Design and characterization of a TrkB-T1-ICD mock peptide: intracellular localization and viability effects in cortical neurons and astrocytes

To establish the specific contribution of TrkB-T1-ICD to excitotoxicity, we generated a CPP mimicking this TrkB-T1 RIP fragment. As the precise TrkB-T1-ICD N-ter amino acid remains unknown (Fig. 1A), we decided to exclude TrkB-T1-ICD TM sequences from the designed peptide, which was designated as Bio-LTT1_Ct_. Thus, this peptide contained a HIV-1 Tat short domain, which allows attached cargoes crossing the BBB and plasma membranes (48, 49), followed by the TrkB-T1 intracellular sequence (Fig. 3A). To facilitate visualization, Bio-LTT1_Ct_ also contained a biotin molecule at the N-ter. A similar peptide containing unrelated sequences corresponding to c-Myc (Bio-TMyc) was used as a negative control. In previous studies, we demonstrated that Bio-TMyc could enter both cortical neurons and astrocytes present in primary cultures (28), but had no effect on neuronal viability either in basal or excitotoxic conditions (39, 50). Immunofluorescence analysis performed in primary cultures of cortical neurons (Fig. 3B) and astrocytes (Fig. 3E) demonstrated peptide entry since early times of treatment (30 min). We also analyzed TrkB-T1 cell distribution by using the isoform-specific antibody mentioned above, although labeling in Bio-LTT1_Ct_-treated cells corresponds to both the treatment peptide and the native complete protein. We observed that Bio-TMyc and Bio-LTT1_Ct_ displayed a different distribution in both cell types. Particularly, Bio-LTT1_Ct_ resembled TrkB-T1 particulate dissemination, suggesting that the TrkB-T1-derived peptide might be located near its corresponding receptor. In addition, a Bio-LTT1_Ct_ fraction was also detected in the nuclei (white arrowheads) of both neurons (Fig. 3B) and astrocytes (Fig. 3E). A cell fractionation assay followed by immunoblot confirmed Bio-LTT1_Ct_ entry into both cortical neurons (Fig. 3C) and astrocytes (Fig. 3F) and a partial migration to the cell nuclei, similar to results obtained with TrkB-T1-ICD. We next investigated the possible effect of this TrkB-T1-derived peptide on cell viability. In neurons (Fig. 3D), Bio-LTT1_Ct_ (25 µM) was not toxic at early incubation times, but decreased the total cell viability (neurons + astrocytes) to 50 ± 11% after 4 h of treatment, a value significantly lower to that obtained with the control peptide Bio-TMyc (109 ± 3%, \**P* < 0.05). This Bio-LTT1_Ct_ effect on cell viability was dose-dependent and not observed for peptide concentrations < 5 µM (data not shown). Cell toxicity related to Bio-LTT1_Ct_ incubation was also detected in cultured astrocytes, where similar effects to those described in neurons were reached at earlier incubation times (2 h, \**P* < 0.05; Fig. 3G). Together, these results suggest that Bio-LTT1_Ct_ could be partially mimicking TrkB-T1-ICD location and effects at least in neurons.

**Fig. 3.**
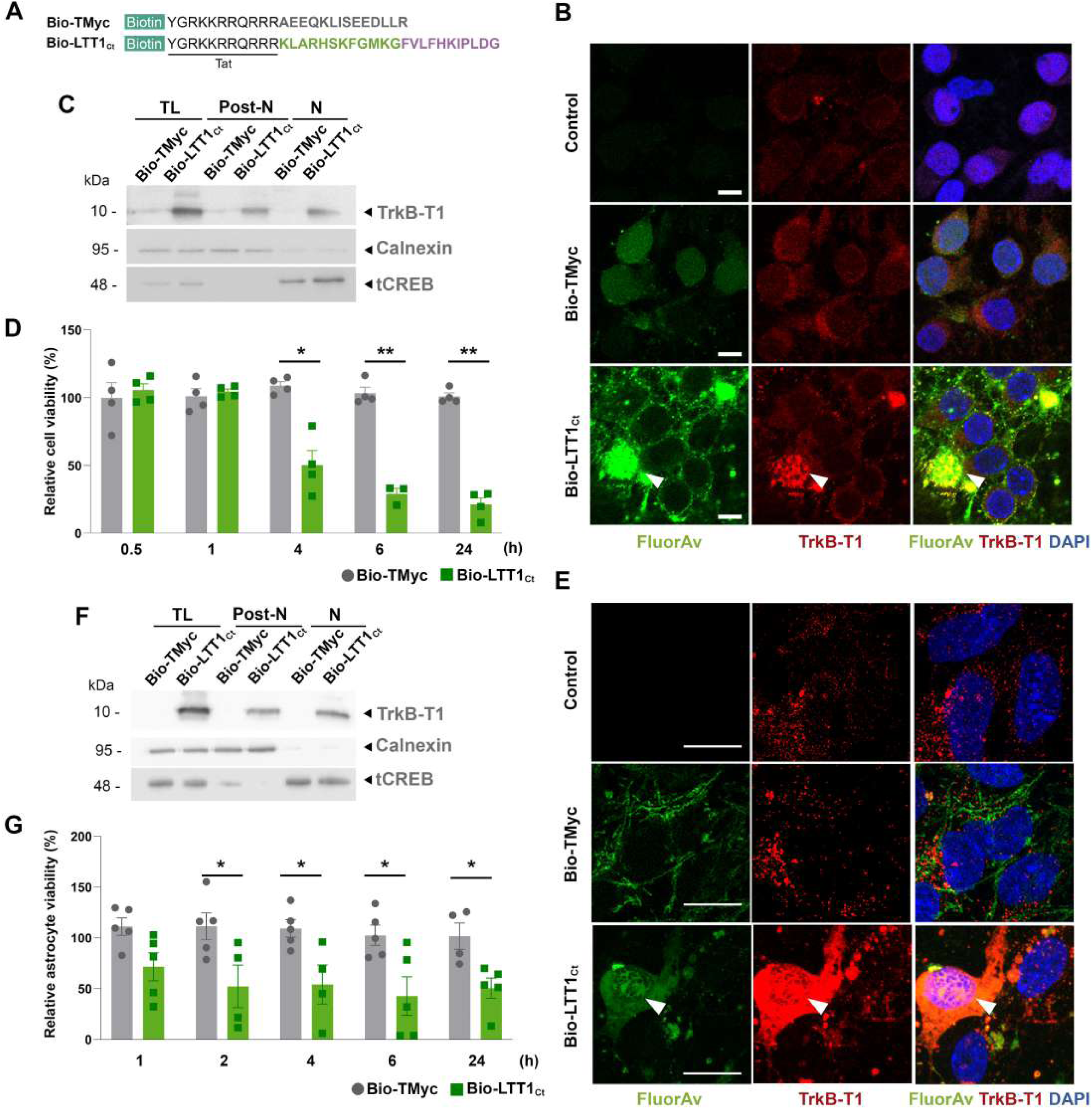
Peptide Bio-LTT1_Ct_ induces cytotoxicity and presents a subcellular distribution similar to endogenous TrkB-T1-ICD. **A** TrkB-T1-derived CPP Bio-LTT1_Ct_ and control peptide Bio-TMyc share a biotin group at the N-ter, followed by the indicated Tat sequence. After that, Bio-TMyc contains amino acids 408 to 421 of the c-Myc protein (grey), and Bio-LTT1_Ct_ includes TrkB-T1 intracellular amino acids 454-476. Some of these amino acids are common to other TrkB isoforms (amino acids 454-465, green), and others are isoform-specific (amino acids 466-476, pink). **B, E** Immunocytochemistry assays of primary neurons (**B**) and astrocytes (**E**) treated with Bio-LTT1_Ct_ or Bio-TMyc (25 µM) for 30 min or left untreated (control). TrkB-T1 and Bio-LTT1_Ct_ distribution was examined by labeling with the isoform-specific antibody (red). Peptides were visualized with Fluorescein Avidin (FluorAv, green). Nuclear location of Bio-LTT1_Ct_ is indicated with white arrowheads. Representative confocal microscopy images are presented. Scale bars represent 10 µm. **C, F** Subcellular fractionation of primary neurons (**C**) and astrocytes (**F**) treated with Bio-LTT1_Ct_ or Bio-TMyc (25 µM) for 30 min. Calnexin was used as a Post-nuclear (Post-N) marker and tCREB as a marker of the nuclear (N) phase. Bio-LTT1_Ct_ was detected using a TrkB-T1 isoform-specific antibody. **D, G** Cell viability of primary neurons (**D**) and astrocytes (**G**) treated with Bio-LTT1_Ct_ or Bio-TMyc (25 µM) for selected times. Viability values are expressed relative to the viability obtained in cells treated with Bio-TMyc for 30 min (considered as 100%). Individual values are presented and data were expressed as the mean ± SEM. Statistical analysis was performed by two-way ANOVA test followed by *post hoc* Tukey’s test (**D**, N = 4) or Sídak’s test (**G**, N = 4-5). Technical replicates were included in each independent experiment (n = 3).

### Characterization of Bio-LTT1_Ct_ effects on the transcription of selected genes in primary cultures of cortical neurons

The presence of Bio-LTT1_Ct_ in the nucleus of neurons and its effects on neuronal viability suggested that this peptide might mimic some of the transcriptional changes induced by excitotoxicity. We have previously shown that overactivation of NMDARs induces a progressive decrease of the promoter activity of CREB and MEF2 (39), TFs which are central to neuronal survival induced by neurotrophins (13, 14) and synaptic activity (51, 52). We then performed reporter assays using promoters with minimal CREB or MEF2 responsive elements, respectively pCRE or pMEF2 (19, 53), controlling the expression of a reporter luciferase gene. Cultures of neurons transfected with these reporter plasmids were treated after 24 h with peptides Bio-LTT1_Ct_ or Bio-TMyc (25 µM, 2.5 h) or left untreated. In pCRE-transfected cells (Fig. 4A, left panel), incubation with Bio-LTT1_Ct_ decreased luciferase activity to 59 ± 6% relative to values obtained in untreated cells, being also significantly inferior to that observed in the presence of the control peptide Bio-TMyc (105 ± 6%, *** *P* < 0.001), which did not affect CREB promoter activity as expected (28, 39). Similar results were obtained for pMEF2-transfected cells (66 ± 8% for Bio-LTT1_Ct_ vs. 98 ± 10% for Bio-TMyc; \**P* < 0.05) (Fig. 4A, central panel). This effect was proved specific since it was reverted by using pMEFmutant, a plasmid similar to pMEF but with inactivating mutations in MEF2 responsive elements (Fig. 4A, right panel). Interestingly, this decrease in promoter activity induced by Bio-LTT1_Ct_ was similar to that induced by 2 h of treatment with NMDA, independently of a 30 min preincubation with Bio-LTT1_Ct_ or Bio-TMyc (Fig. 4A, columns labeled as +). These results suggest that the TrkB-T1 intracellular sequence plays by itself a fundamental role in the transcriptional regulation of CREB/MEF2-dependent genes during excitotoxicity.

**Fig. 4.**
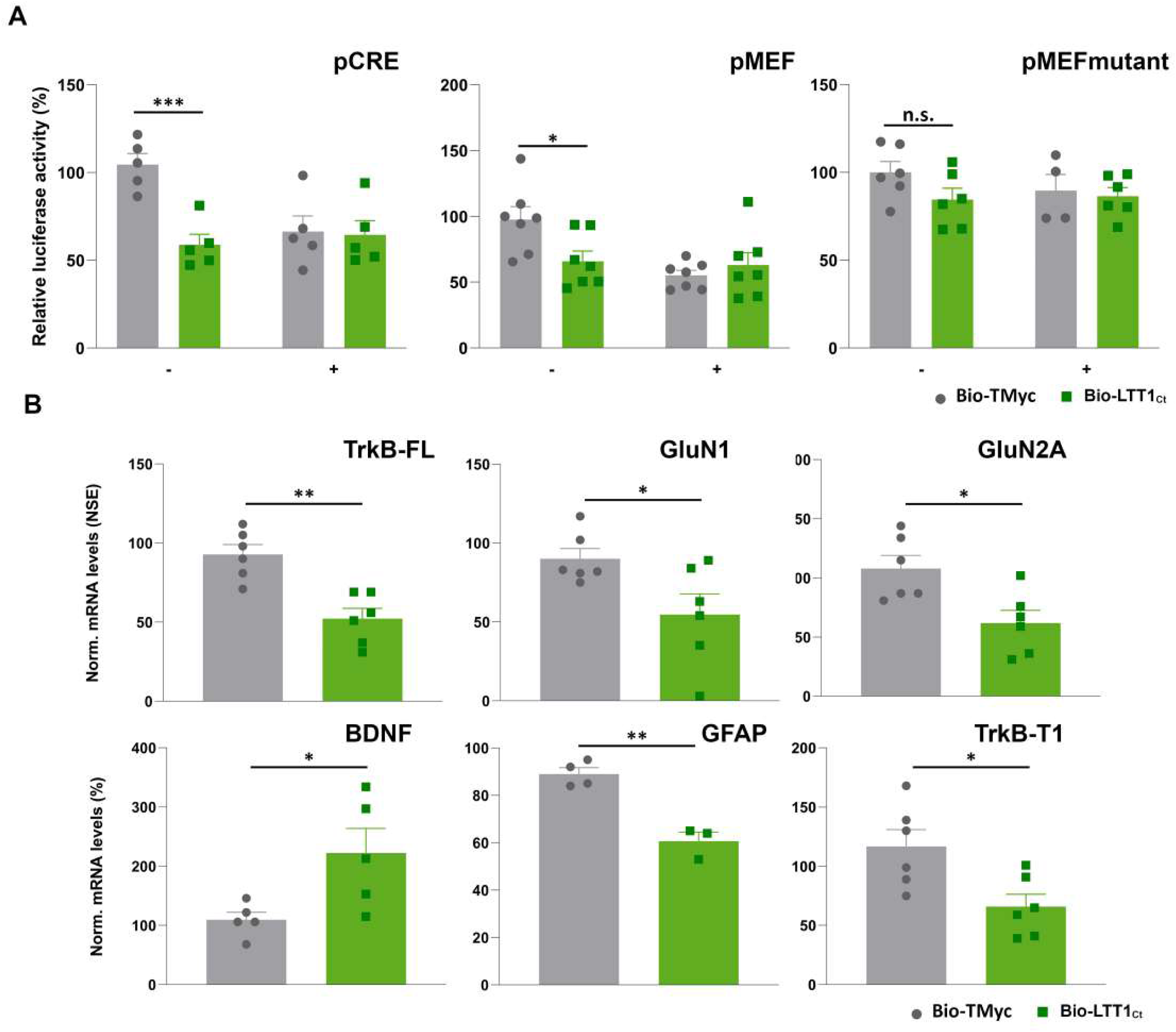
Bio-LTT1_Ct_ effects on transcriptional regulation leads to an excitotoxicity-like scenario. **A** Effect of Bio-LTT1_Ct_ treatment on CRE and MEF2 promoter activities. Cultures transfected with luciferase reporter plasmids pCRE, pMEF and pMEFmut were treated for 2 h with Bio-LTT1_Ct_ or Bio-TMyc (25 µM) (-conditions). Alternatively, cultures were preincubated with the peptides for 30 min and then subjected to excitotoxicity with NMDA for 2 h, CPPs remaining in the media (+ conditions). Individual data are presented and expressed as the mean ± SEM of luciferase activities and presented relative to those obtained in untreated cultures. Data were analyzed by two-way ANOVA test followed by *post hoc* Sídak’s test (N = 5-7, n = 4)**. B** Effect of Bio-LTT1_Ct_ treatment on mRNA levels of CREB/MEF2-regulated genes. Cultures were treated for 2 h with Bio-LTT1_Ct_, Bio-TMyc (25 µM) or left untreated. Levels of mRNA were normalized to neuronal specific enolase (NSE) (genes mainly expressed in neurons: TrkB-FL, GluN1 and GluN2A) or GAPDH (genes expressed both in neuronal and glial cells: BDNF, GFAP and TrkB-T1). Individual data are presented and expressed as the mean ± SEM of mRNA levels relative to values found in untreated cultures. Data were analyzed by Student’s *t*-test (N = 4-6, n = 3).

We next analyzed mRNA levels of several downstream genes to investigate if Bio-LTT1_Ct_ also mimicked transcriptional changes induced by excitotoxicity (39, 54). Primary cultures of neurons were treated for 2 h with Bio-LTT1_Ct_, Bio-TMyc or left untreated (Fig. 4B). Compared to Bio-TMyc cultures, Bio-LTT1_Ct_ incubation significantly reduced the relative levels of pro-survival genes such as the *Ntrk2* splicing form encoding TrkB-FL or the GluN1 and GluN2A NMDAR subunits, while it increased those of BDNF mRNA. The changes induced by Bio-LTT1_Ct_ were similar to those described in cultures subjected to excitotoxicity by NMDA incubation (39, 54). However, in contrast to results obtained in excitotoxic conditions (29, 55), levels of characteristic astrocytic mRNAs encoding GFAP and TrkB-T1 were significantly reduced by Bio-LTT1_Ct_. These results demonstrate that, concurrent to nuclear migration, Bio-LTT1_Ct_ alters CREB and MEF promoter activities as well as the expression of some important neuronal genes related to survival/death choices, reproducing an excitotoxic cell environment with some peculiarities that merit further characterization. Moreover, they strongly suggest that the RIP-product TrkB-T1-ICD might be involved in the control of gene transcription and have a relevant role in excitotoxicity.

### Characterization of Bio-LTT1_Ct_ effects on mice brain cortex after intranasal peptide administration

We next examined if, by disturbing the function of neurons and/or glial cells, Bio-LTT1_Ct_ was able to promote changes in mice brain mimicking some stroke pathological mechanisms. We used intranasal administration, a non-invasive delivery route to the brain parenchyma that takes advantage of the trigeminal and olfactory pathways and the microvasculature to bypass the BBB (56). Particularly, the administration of CPPs by this route is currently under intense investigation since it provides further feasibility to naso-brain drug delivery (57, 58). By these means, mice were treated with Bio-LTT1_Ct_ or Bio-TMyc (2.5 nmol/g) CPPs for 1 or 48 h, using the vehicle as an additional control (Fig. 5A). First, we approached the analysis of brain delivery and stability of the biotinylated peptides by wide-field imaging, using Cy5-Streptavidin (Cy5-Strep, magenta) for CPP labeling combined with NeuN (green) (Fig. 5B). One hour after administration, animals treated with Bio-TMyc or Bio-LTT1_Ct_ showed peptide delivery mainly into the cortical brain area. Moreover, these peptides could be detected in both brain hemispheres, independently of the specific nostril of application. Furthermore, the presence of both CPPs in the brain cortex decreased to background levels in mice sacrificed 48 h after administration. Using confocal microscopy, we next investigated in more detail the distribution of Bio-TMyc and Bio-LTT1_Ct_ in the brain (cortical and subcortical areas) of animals treated for 1 h with these CPPs (Fig. 5C, D). Compared to results obtained in animals treated with vehicle, Bio-TMyc and Bio-LTT1_Ct_ were detected in most cortical cells, distributing in cell bodies and projections of NeuN^+^ (asterisks) and NeuN^-^ cells (arrowheads) (Fig. 5C). In contrast, these biotin-labeled peptides could not be detected in the subcortical areas of these same coronal sections (Fig. 5D). Above results demonstrate an efficient nose-to-brain delivery of Bio-LTT1_Ct_ and Bio-TMyc peptides, able to cross the BBB of undamaged animals and reach the cerebral cortex. In addition, the limited stability of these peptides in the tissue provides a temporal frame of action to analyze peptide distribution and/or their brain effects.

**Fig. 5.**
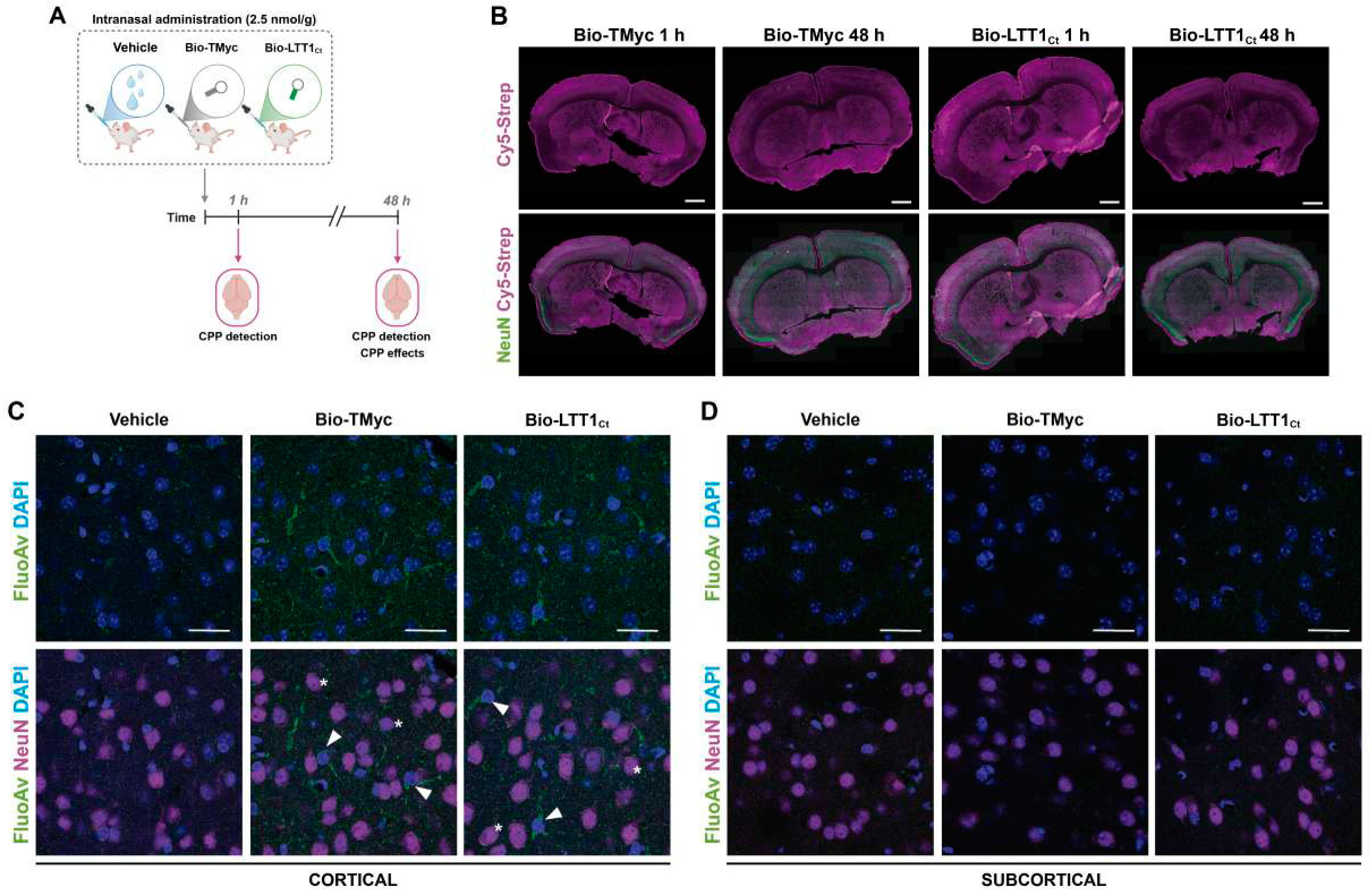
Intranasal (*i.n.*) administration of Bio-LTT1_Ct_ leads to effective brain delivery. **A** Experimental design. Adult male mice receiving a single dose of vehicle, Bio-TMyc or Bio-LTT1_Ct_ (2.5 nmol/g) by *i.n.* administration were examined 1 or 48 h later, after intracardiac perfusion with 4% PFA. Peptide levels were studied both at 1 and 48 h after treatment while the effect of CPPs on brain homeostasis was analyzed at 48 h. **B** Peptides were visualized with Cy5-Streptavidin (Cy5-Strep, magenta) in fixed coronal sections of animals treated with Bio-TMyc or Bio-LTT1_Ct_ for 1 or 48 h, as indicated. Neurons were also identified using the NeuN antibody (green). Wide-field images were obtained with a Cell Observer Microscope. **C** and **D** Detailed analysis of Bio-LTT1_Ct_ delivery to the mice cortex after 1 h of CPP administration. Bio-LTT1_Ct_ and Bio-TMyc were detected in coronal sections by Fluorescein Avidin D (FluorAv, green), neurons with NeuN (magenta) and cell nuclei with DAPI (blue). Peptide delivery was observed in neuronal (asterisks) and non-neuronal cells (arrowheads). Representative confocal microscopy images correspond to single sections from cortical and subcortical areas. Scale bars represent 20 μm (N = 4).

In previous work, we demonstrated that ischemic stroke induced an early decrease of TrkB-FL levels in neurons of the infarcted area accompanied by an increase of TrkB-T1 (29), together with increased expression of TrkB-T1, GFAP and C3 in astrocytes of the emerging fibroglial scar (28). We hypothesized that these actions could be promoted by the TrkB-T1-ICD formed as a result of TrkB-T1 RIP, induced by *in vivo* excitotoxicity secondary to stroke. To prove this hypothesis, we first analyzed the effect of Bio-LTT1_Ct_ treatment on the cortical levels of TrkB isoforms 48 h after peptide administration (Fig. 6A, B). Double immunohistochemistry with NeuN showed that, compared to animals treated with Bio-TMyc, Bio-LTT1_Ct_ induced an increase of TrkB-T1 (Fig. 6A) and a decrease of TrkB-FL (Fig. 6B) that seemed to affect both cortical neurons and astrocytes. The analysis of the Bio-LTT1_Ct_ effects on TrkB-T1 levels was possible because, as shown before, this peptide becomes undetectable in the brain cortex at this time-point (48 h, see Fig. 5B) and, therefore, it does not interfere with the detection of the native peptide using the same specific antibody. Concurrent to TrkB-T1 upregulation, levels of GFAP also increased in the cortical astrocytes of animals treated with Bio-LTT1_Ct_ (Fig. 6C). Moreover, this peptide not only promoted astrocyte reactivity but also their pro-inflammatory profile, these astrocytes being co-stained with GFAP and C3 antibodies (Fig. 6D). Altogether, these results demonstrate that, in non-ischemic animals, Bio-LTT1_Ct_ can promote by itself some stroke characteristic effects regarding the expression of specific proteins in neurons and astrocytes.

**Fig. 6.**
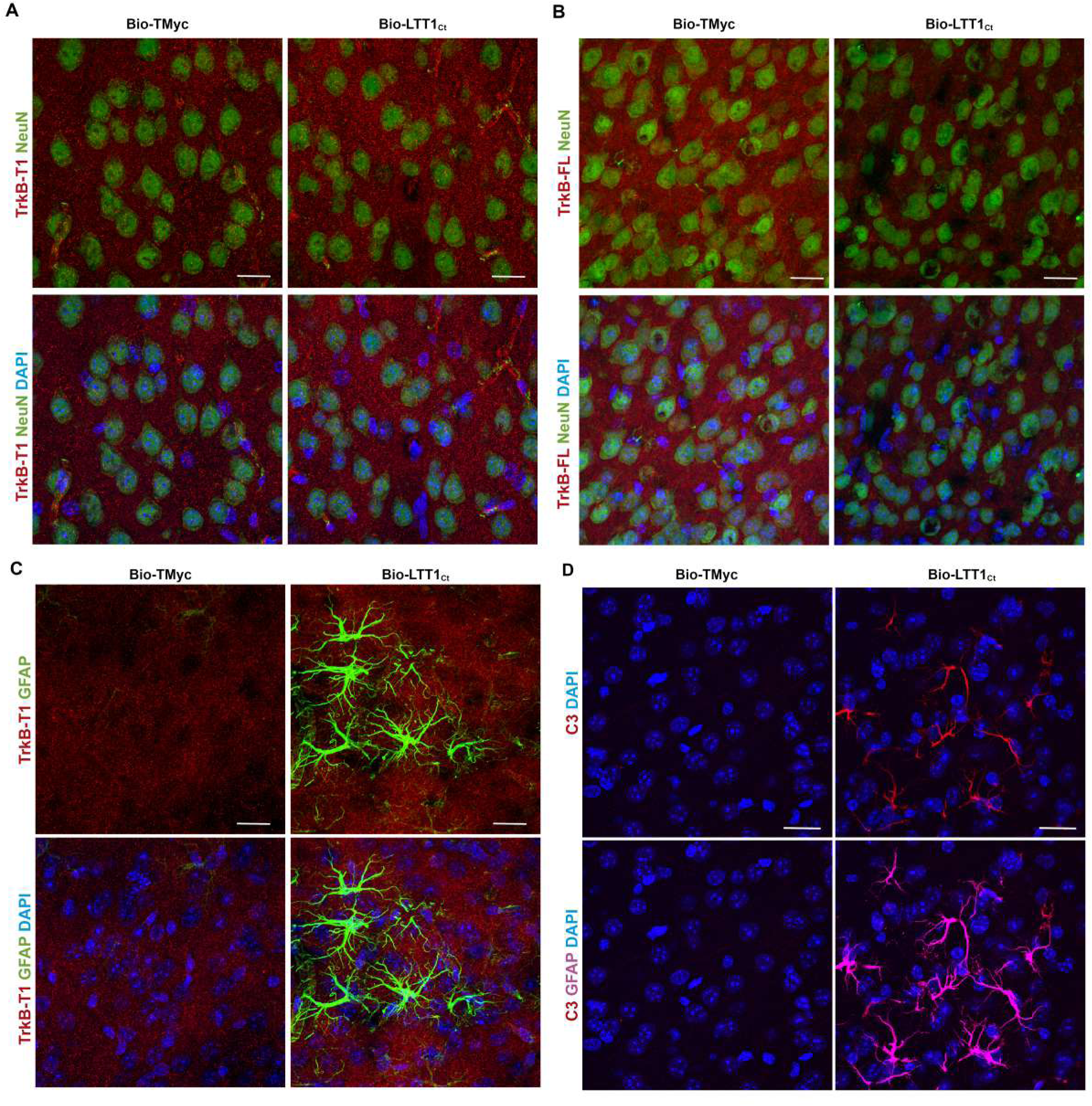
Bio-LTT1_Ct_ induces stroke-like changes in neurons and astrocytes after intranasal administration. **A, B** Double immunohistochemistry of TrkB-T1 (**A**) and TrkB-FL (**B**) in male mice cortex 48 h after CPP administration. Both TrkB isoforms were detected with isoform-specific antibodies (red), counterstained with NeuN (green) for neurons and DAPI (blue) for nuclei staining. **C** Double immunohistochemistry of TrkB-T1 (red) and GFAP (green) in male mice cortex 48 h after CPP administration. DAPI (blue) for nuclei staining is also shown. **D**. Double immunohistochemistry of complement protein C3d (red) and GFAP (magenta) in male mice cortex 48 h after CPP administration. DAPI (blue) for nuclei staining is also shown. Representative confocal microscopy images of cortical areas correspond to maximum intensity projections. Scale bars represent 10 μm (**A and B**) and 20 μm (**C and D**) (N = 4).

Finally, we explored a possible effect of Bio-LTT1_Ct_ administration on microglia activation (Fig. 7). In the ischemic brain, microglia display a broad variety of morphological configurations not limited to ramified or amoeboid cell shapes, corresponding respectively to surveilling or activated microglia (59, 60). A preliminary observation of Iba1 expression in the cortical region of animals treated with Bio-LTT1_Ct_ for 48 h revealed subtle but detectable morphological differences in the microglia, compared to vehicle or Bio-TMyc treated animals, together with a strong increase in Iba1 expression (Fig. 7A for details and Fig. S3 for complete tiles). To characterize in depth these differences, we next performed a tridimensional reconstruction using the cortical tiles acquired after Iba1 staining, which was followed by cell segmentation and measurement of different morphological parameters (*i.e.,* volume, sphericity, area, and shortest ellipsoid), count of microglia cells, and mean Iba1 intensity using AIVIA software (Fig. 7B, C). Comparison of representative 3D reconstructed cells showed more rounded and ameboid-like morphological characteristics for microglia in Bio-LTT1_Ct_ treated animals (Fig. 7B) which correlated with a significant increase of sphericity and decreases of volume and area compared to Bio-TMyc animals (Fig. 7C). Additionally, these experiments also revealed that Bio-LTT1_Ct_ administration induced an increase in the number of microglial cells and mean Iba1 intensity (Fig. 7C). In contrast, treatment with the control peptide Bio-TMyc did not have any effect on microglial morphology or count, the results being very similar to those obtained with the vehicle (Fig. 7B, C and Fig. S3). Furthermore, a Principal Component Analysis (PCA) performed using previous parameters, showed a clear separation between the Bio-LTT1_Ct_ experimental group and the other two groups analyzed, indicating a clear effect of this peptide in cortical microglia (Fig. 7D). Then, we performed Uniform Manifold Approximation (UMAP) analysis of all Iba1^+^ cells to analyze the distribution of the microglia from the differently treated animals according to the parameters mentioned above (Fig. 7E). Again, cells from animals treated with Bio-LTT1_Ct_ (Fig 7F) showed an opposite behavior to those from animals treated with Bio-TMyc (Fig 7G) or vehicle (Fig. S4). To deepen in the characterization of the morphological subtypes, we used unsupervised clustering using the K-means algorithm to evaluate the presence of different morphological clusters according to our measurements. This analysis allowed us to differentiate five different groups, represented in the UMAP plot together with their corresponding 3D reconstructions (Fig. 7H). This result is not unexpected since, as mentioned before, microglia are a highly heterogeneous and dynamic cell population and, thus, not all cells in the cortex will present the same morphological patterns. A correlation analysis confirmed that Bio-LTT1_Ct_ administration induced an increase of microglia having a more rounded and activated-like morphology, while promoted a decrease in the levels of the resting-like groups (Fig 7I). As before, the control groups (vehicle and Bio-TMyc) presented an opposite morphological pattern, being enriched in resting-like microglia. Altogether, these changes clearly indicate that Bio-LTT1_Ct_ administration is able to promote by itself an activation state of the microglia which is comparable to that taking place in stroke (61, 62). We also conclude that TrkB-T1-ICD is sufficient to trigger transcriptional changes in neurons and glial cells after stroke, depleting the expression of pro-survival proteins and promoting astrocyte and microglia reactivity, hallmarks of ischemic stroke and main contributors to brain damage, which can be mimicked by Bio-LTT1_Ct_ treatment in the absence of stroke.

**Fig. 7.**
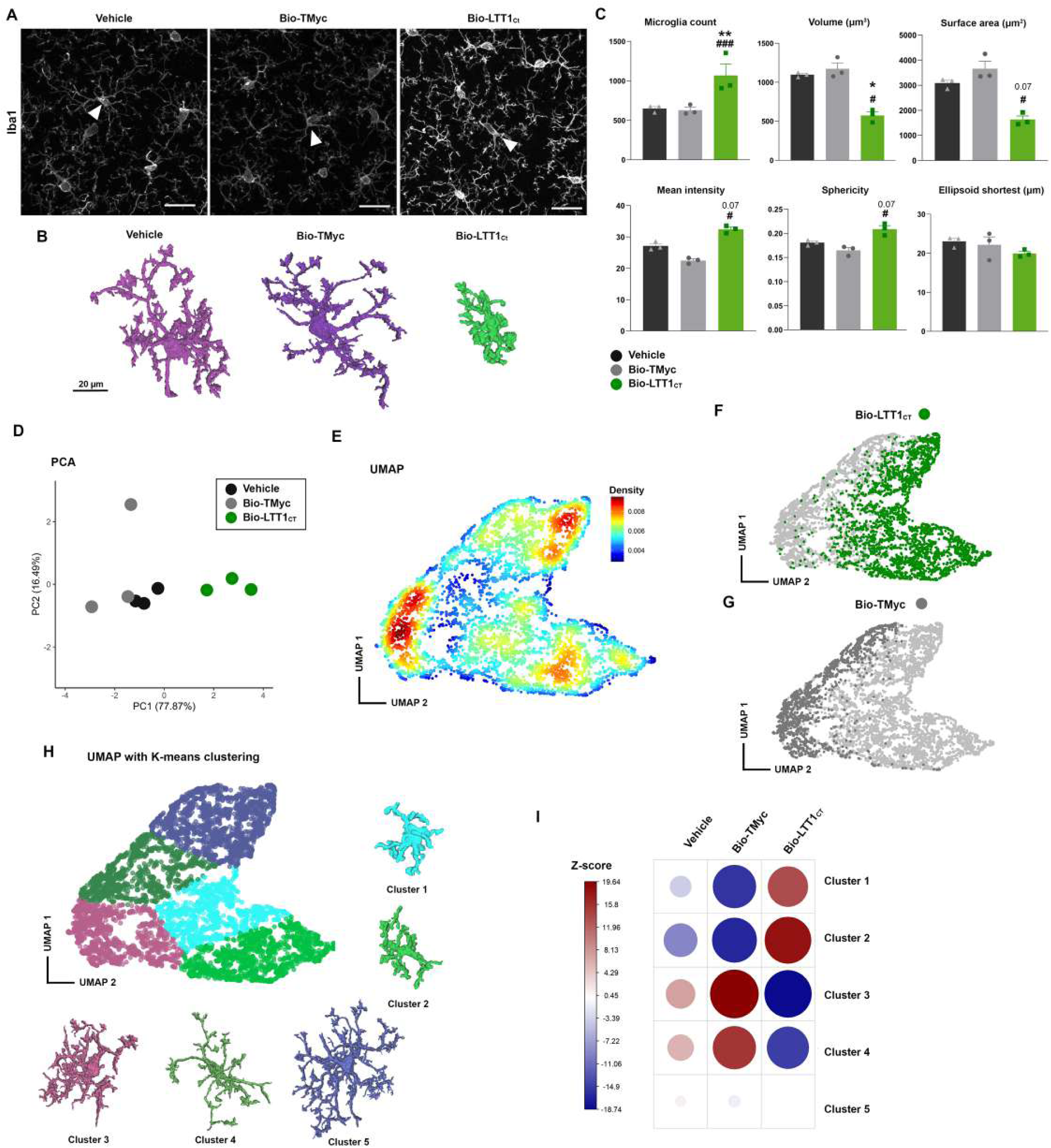
Bio-LTT1_Ct_ promotes microglia activation after intranasal administration. **A.** Immunohistochemistry of Iba1 after 48 h of peptide treatment. Representative images from selected areas of tiles obtained by confocal microscopy from cortical areas are shown (see Fig. S3). Images correspond to maximum intensity projections. Scale bars represent 20 μm. **B.** 3D representation of selected cells from the images shown in A. Morphological reconstruction was obtained from 3D segmentation of each Iba1 positive cell in the original images. **C**. Bar plots represent the individual mean value of selected morphological parameters and number of microglial cells for each mouse and treatment. Statistical analysis was carried out by using two-way ANOVA test followed by Tukey’s multiple comparison test (* for Vehicle *vs* Bio-LTT1_Ct_, # for Bio-TMyc *vs* Bio-LTT1_Ct_). Data are expressed as the mean ± SEM (N = 3). **D**. PCA representation based on the parameters shown in C for animals treated with vehicle, Bio-TMyc or Bio-LTT1_Ct_. Variance represented by the First (PC1) and Second (PC2) Principal Components are shown. **E.** UMAP-density plot representation of microglial cells from all mice and groups of treatment. Cell density is presented using UMAP1 and UMAP2 dimensions. **F and G.** Cells corresponding to Bio-LTT1_Ct_ (green) or Bio-TMyc (grey) treated animals are respectively highlighted. **H.** UMAP representation of unsupervised K-means clustering based on previous parameters. Clusters detected are represented in the same colors as their representative morphological reconstruction. **I.** Correlation plot between treatment conditions and presence of the identified morphological clusters. Z-score is presented, red indicating enrichment and blue indicating depletion.

## Discussion

The results presented in this work support the hypothesis that the truncated neurotrophin receptor, TrkB-T1, plays a key role in stroke pathology mainly mediated by its RIP product, TrkB-T1-ICD. This proposition is based on the demonstration that the short TrkB-T1 intracellular sequence is able to recapitulate fundamental aspects of stroke related to neurotoxicity, glial reactivity and neuroinflammation. These observations complement current understanding about the different excitotoxicity-induced mechanisms that, by acting on the TrkB isoforms, contribute to impaired BDNF-signaling. Particularly, NMDAR overactivation inverts the transcriptional pattern of *Ntrk2* gene in neurons, favoring the expression of TrkB-T1 over TrkB-FL (29). Moreover, TrkB-FL transcriptional depletion is complemented by retrograde transport of the encoded protein to the Golgi and calpain-processing (29, 38–40), producing a truncated membrane receptor similar to TrkB-T1, which might act as an additional dominant-negative receptor, and a cytosolic fragment containing the TK domain, which translocated to the nucleus (63). Regarding TrkB-T1, assessment of its contribution to stroke pathology has proven to be far more complex. The increase of TrkB-T1 in excitotoxic neurons, where it inhibits BDNF/TrkB-FL-signaling (21), significantly contributes to neurotoxicity (29). However, there are other TrkB-FL-independent mechanisms exerted by TrkB-T1 in neurons and astrocytes that also participate in stroke pathologic features. Firstly, we recently demonstrated that TrkB-T1 accumulates with GFAP in activated pro-inflammatory astrocytes at the infarct border from early times of ischemia (28). This pro-inflammatory activation is originated by protein interactions established by the TrkB-T1 isoform-specific sequence under excitotoxic conditions, likewise modulating brain infiltration of immunoglobulins and macrophages, and activation of the microglia occurring in the infarct core and adjacent tissue (28). Secondly, astrocytic TrkB-T1 upregulation in ischemia is also associated with the promotion of cerebral edema and exacerbated brain damage (33). This is important because cerebral edema is an acute complication of ischemic stroke often associated with poor prognosis (64). In previous work, we proposed that, in addition to TrkB-T1 upregulation, the isoform fragments generated by RIP might strongly contribute to stroke pathology (38). Indeed, we demonstrated that TrkB-T1 shedding releases a receptor ectodomain that contributes to reduced neurotrophic support in neurons. Herein, we present novel and robust data demonstrating that the TrkB-T1-ICD fragment, produced by γ-secretase intramembrane cleavage of a membrane-tethered C-ter fragment, partly migrates from the cytoplasm to the nucleus, where it probably affects the transcription of some important genes for survival/death choices, as well as neuronal morphology and function.

While the contribution of TrkB-T1 RIP to stroke pathology has been strongly established in neurons, some controversy remains regarding astrocytes. Treatment of astrocyte-enriched cultures with NMDA induces intracellular cascades leading to TrkB-T1 and GFAP upregulation, in agreement with data showing that, despite low levels of NMDARs, they are able to mediate Ca^2+^-waves regulating important intracellular processes such as gene expression (47, 65). However, differently from cultured neurons, NMDA does not induce TrkB-T1 RIP in our model of cultured astrocytes. This seems contradictory to reported results showing increased activity of brain γ-secretase early after stroke and, particularly, worsened brain damage and functional outcome due to Notch-1 RIP (66, 67). Notably, Notch-1-signaling is involved in the regulation of the proliferation of reactive astrocytes in the peri-infarct area after stroke (68). Therefore, in contrast to results obtained *in vitro,* TrkB-T1 RIP is probably induced by ischemia in astrocytes *in vivo* and, similarly to Notch-ICD, TrkB-T1-ICD might also contribute to regulate astrocyte reactivity. The different response of astrocytes *in vitro* and *in vivo* is not surprising because it has been previously recognized that conventional 2D astrocytes cultures present important limitations to reproduce the microenvironment of these cells in the brain (69). However, alternative possibilities should be also considered. For example, TrkB-T1-ICD might be only produced in excitotoxic neurons and transmitted to glial cells through extracellular vesicles (EVs). These vesicles are essential mediators of brain intercellular communication able to transport signaling molecules such as RNAs, proteins and lipids. Moreover, they can be released by neurons, astrocytes, oligodendrocytes, and other CNS cells, and play key roles in physiological processes (*e.g.,* synaptic plasticity, brain development). Additionally, they are also involved in pathological mechanisms and, in the case of NDDs, they contribute to the spreading of proteins such as amyloid β, tau, ɑ-synuclein, prions, and TDP-43, aggravating neurodegeneration and accelerating disease progression (70). In stroke, neuron-derived EVs carry a variety of miRNAs that affect microglia by distinct mechanisms and influence stroke progression (71). However, the crosstalk between neurons and astrocytes via EVs in stroke has been limitedly explored. Therefore, it is of high interest and will merit further characterization to establish if TrkB-T1-ICD could be an EVs cargo contributing to pro-inflammatory responses after stroke through EV-mediated communication among different brain cell types.

To complete the elucidation of the specific contribution of TrkB-T1-ICD to stroke, in this work we developed a brain-accessible CPP, Bio-LTT1_Ct_, containing the complete intracellular TrkB-T1 region. The use of CPPs is attracting great interest due to their novelty and useful properties, including their capacity to translocate across the BBB, ability to penetrate into many different cell types, and low toxicity (72, 73). Indeed, these peptides are not only useful as non-invasive delivery systems of therapeutic compounds but, as shown here, as remarkable tools to approach the mechanisms of disease, particularly those affecting the CNS. From early times of treatment, Bio-LTT1_Ct_ distributes in cultured neurons and astrocytes and mice cortex, promoting some of the responses that excitotoxicity and stroke have in those specific cells. Thus, Bio-LTT1_Ct_ partial migration to the nucleus in primary cultures resembles that of TrkB-T1-ICD induced by NMDA. However, differently from excitotoxic conditions, which do not decrease the viability of the small astrocyte subpopulation present in neuronal-enriched cultures (45, 46), Bio-LTT1_Ct_ is toxic for both neurons and astrocytes. In contrast, this peptide does not seem to be lethal *in vivo,* at least 48 h after brain administration, establishing a differential response of astrocyte *in vitro* and *in vivo* that might be due again to technical limitations of these conventional 2D cell cultures (69). Parallel to nuclear migration and before cell death, Bio-LTT1_Ct_ also promotes transcriptional changes in neuronal-enriched cultures similar to those induced by NMDA treatment. For instance, this peptide decreases the promoter activity of the pro-survival TFs CREB and MEF2 in a similar way than excitotoxicity does and, furthermore, the combined treatment with NMDA does not further aggravate the results. Based on these findings, we conclude that TrkB-T1 intracellular region has a critical role in CREB and MEF2 activities and their downregulation after excitotoxic injury. Accordingly, a brief treatment with Bio-LTT1_Ct_ (2 h) alters the mRNA levels of important neuronal genes related to survival/death (*i.e.,* TrkB-FL, NMDAR-subunits and BDNF), reproducing changes characteristic of excitotoxicity (39, 54). Only TrkB-T1 and GFAP mRNA levels do not correspond to those expected for excitotoxicity, suggesting that astrocytes present in mixed cultures might be already deeply affected by Bio-LTT1_Ct_ at this time point. In contrast, experiments performed *in vivo* show that Bio-LTT1_Ct_ reproduces in the cortex relevant changes in protein levels induced by ischemia such as the imbalance of the TrkB isoforms and the upregulation of GFAP and C3 expression in astrocytes. Finally, Bio-LTT1_Ct_ also promotes important morphological changes in the microglia resembling the activation pattern typically induced in the first hours after brain ischemia (28). These results strongly suggest that TrkB-T1-ICD is involved in the regulation of gene transcription *in vivo*, affecting neurons, astrocytes and microglia. A limitation of these studies could be that Bio-LTT1_Ct_ does not include the complete TrkB-T1-ICD sequence, as the precise amino acid cleaved by γ-secretase in the TrkB-T1 TM region remains unknown and, despite having a similar behavior, we could be missing some effects of the native TrkB-T1-ICD. Nonetheless, our results support TrkB-T1 as a master protein controlling complex interactions among excitotoxicity, neurotrophic-signaling, glial reactivity, and neuroinflammation, all processes central to stroke pathology. In addition, TrkB-T1-ICD, resulting from excitotoxicity-induced TrkB-T1 RIP, emerges as a main mediator of these TrkB-T1 pathological actions.

### Conclusions

A deep characterization of the interrelation among the different pathological processes participating in ischemic injury is pivotal for the development of brain-accessible therapeutic molecules able to provide multimodal and multicellular brain effects and, therefore, become integrative stroke protection strategies. In line with our previous findings, herein we show that the neurotrophin receptor TrkB-T1 —and particularly a small fragment containing its intracellular region, TrkB-T1-ICD, product of excitotoxicity-induced RIP— has critical roles in neurotoxicity, aberrant neurotrophic-signaling, glial reactivity, and neuroinflammation. This observation is of pivotal importance because these relevant pathological signatures are not only linked to stroke but also to many other acute and chronic CNS disorders similarly associated with excitotoxicity. The sole addition of Bio-LTT1_Ct_, a brain-accessible peptide mimicking this intracellular TrkB-T1 sequence, is able to recapitulate stroke-related effects both *in vitro* and *in vivo*, with a particular effect on neurotoxicity and the reactivity of the glial and microglial subpopulations. Altogether, this study demonstrates for the first time the importance of TrkB-T1-ICD as a key modulator of glial reactivity and neuroinflammation in acute CNS pathologies and proposes an important target for neuronal and glial cells protection under excitotoxic conditions.

## Abbreviations

APMA: p-aminophenylmercuric acetate
BBB: blood-brain barrier
BDNF: brain-derived neurotrophic factor
C3: complement component 3
CNS: Central nervous system
CPP: cell-penetrating peptide
CREB: cAMP response element-binding protein
DIVs: days *in vitro*
ECD: extracellular domain
GABA: gamma aminobutyric acid
GFAP: glial fibrillary acidic protein
GO: gene ontology
Iba1: Ionized calcium-binding adapter molecule 1
ICD: intracellular domain
*i.n.*: intranasal administration
MEF2: myocyte enhancer factor 2
MP: metalloproteinase
NDDs: neurodegenerative diseases
NMDARs: N-methyl-D-aspartate receptors
n.s.: non-significant
NSE: neuronal specific enolase
RIP: regulated intramembrane proteolysis
SEM: standard error of the mean
SCI: spinal cord injury
TFs: transcription factors
TK: tyrosine kinase
TM: transmembrane sequence
TrkB: tropomyosin-related kinase B receptor
TrkB-FL: full-length TrkB

## Supplementary information

**Additional file 1: Table S1.** Reagents and resources.

**Additional file 2: Fig. S1.** Subcellular fractionation of cultured astrocytes treated with APMA. Cultures were incubated for 4 or 6 h with APMA (100 µM). TrkB-T1-ICD was detected by the TrkB-T1 isoform-specific antibody. Correct cell fractionation was checked using calnexin as a Post-N marker and tCREB as an N marker.

**Additional file 3: Fig. S2.** Excitotoxicity-induced RIP contributes to neuronal death of primary cultures of cortical neurons. Neuronal viability was measured in cultures treated with NMDA (100 µM) for 1 and 2 h, with or without a 30 min pre-treatment with a broad-spectrum metalloprotease inhibitor (TAPI-2, 10 µM). Individual data of neuronal viability relative to that of untreated cells are presented as the mean ± SEM (N = 3). Technical replicates were included in each independent experiment (n = 3). Data were statistically analyzed using a paired Student’s t-test.

**Additional file 4: Fig. S3.** Microglia morphological analysis. Representative Iba1 and DAPI staining of complete tiles from the cortical area of animals treated with vehicle, Bio-TMyc or Bio-LTT1_Ct_. The fields selected for display in Fig. 7A are indicated with a white square and enlarged next to its corresponding tile. Scale bars are indicated in each case.

**Additional file 5: Fig. S4.** UMAP plot represents microglial cells from all mice and groups of treatment. Cells are presented using UMAP1 and UMAP2 dimensions. **A** and **B** panels highlight cells from the Bio-LTT1_Ct_ treated animals (green) and those from animals treated with Bio-TMyc (grey) or vehicle (black), respectively.

**Additional file 6:** Segmentation code used for analysis.

## Declarations

### Ethics approval and consent to participate

animal procedures were performed in compliance with European Union Directive 2010/63/EU and approved by the CSIC and Comunidad de Madrid (Ref PROEX 276.6/20) ethics committees.

### Consent for publication

not applicable.

### Availability of data and materials

all data generated or analyzed during this study are included in this published article and its supplementary information files.

### Authors’ contributions

LUT designed and performed the experiments, analyzed data, contributed to the original draft, prepared the figures and reviewed/edited the manuscript draft; MCS contributed to experimental design and analysis, supervised project and reviewed/edited the manuscript; MD-G secured funding, designed experiments and supervised project, contributed to formal analysis, wrote the original draft, and reviewed/edited the manuscript. All authors read and approved the final manuscript.

## Acknowledgements

We are grateful to Dr. Sergio Gascón (Instituto Cajal, CSIC) for his initial technical guidance with astrocyte culture, to Candela Chovas for her contributions to the preliminary experiments with astrocytes, Elena Torres Campos for her help with *in vivo* experiments and to members of our group for helpful discussions. We acknowledge the contribution of the Genomics and Microscopy (SEMOC) units (IIBM, CSIC), with an especial mention to Drs. M. Martín Belinchón and B. Acosta for technical support in image acquisition for 3D microglial reconstruction studies. The Advanced Light Microscopy Service at Centro Nacional de Biotecnología (CNB–CSIC) is acknowledged for assistance with the AIVIA software. We are also grateful to “Conexión de Nanomedicina” from CSIC for their support.

## References

1. Collaborators GBDS. Global, regional, and national burden of stroke and its risk factors, 1990-2019: a systematic analysis for the Global Burden of Disease Study 2019. Lancet Neurol. 2021;20:795–820.

2. Xu S, Lu J, Shao A, Zhang JH, Zhang J. Glial Cells: Role of the immune response in ischemic stroke. Front Immunol. 2020;11:294.

3. Shi FD, Yong VW. Neuroinflammation across neurological diseases. Science. 2025;388:eadx0043.

4. Lombardozzi G, Castelli V, Giorgi C, Cimini A, d’Angelo M. Neuroinflammation strokes the brain: a double-edged sword in ischemic stroke. Neural Regen Res. 2026; 21:1715–22.

5. Liddelow SA, Guttenplan KA, Clarke LE, Bennett FC, Bohlen CJ, Schirmer L, et al. Neurotoxic reactive astrocytes are induced by activated microglia. Nature. 2017;541:481–7.

6. Shen XY, Gao ZK, Han Y, Yuan M, Guo YS, Bi X. Activation and role of astrocytes in ischemic stroke. Front Cell Neurosci. 2021;15:755955.

7. Escartin C, Galea E, Lakatos A, O’Callaghan JP, Petzold GC, Serrano-Pozo A, et al. Reactive astrocyte nomenclature, definitions, and future directions. Nat Neurosci. 2021;24:312–25.

8. Hasel P, Rose IVL, Sadick JS, Kim RD, Liddelow SA. Neuroinflammatory astrocyte subtypes in the mouse brain. Nat Neurosci. 2021;24:1475–87.

9. Verkhratsky A, Parpura V, Li B, Scuderi C. Astrocytes: The housekeepers and guardians of the CNS. Adv Neurobiol. 2021;26:21–53.

10. Lei L, Parada LF. Transcriptional regulation of Trk family neurotrophin receptors. Cell Mol Life Sci. 2007;64:522–32.

11. Middlemas DS, Lindberg RA, Hunter T. TrkB, a neural receptor protein-tyrosine kinase: evidence for a full-length and two truncated receptors. Mol Cell Biol. 1991;11:143–53.

12. Niu C, Yue X, An JJ, Bass R, Xu H, Xu B. Genetic dissection of BDNF and TrkB expression in glial cells. Biomolecules. 2024;14:91.

13. Bonni A, Brunet A, West AE, Datta SR, Takasu MA, Greenberg ME. Cell survival promoted by the Ras-MAPK signaling pathway by transcription-dependent and - independent mechanisms. Science. 1999;286:1358–62.

14. Liu L, Cavanaugh JE, Wang Y, Sakagami H, Mao Z, Xia Z. ERK5 activation of MEF2-mediated gene expression plays a critical role in BDNF-promoted survival of developing but not mature cortical neurons. Proc Natl Acad Sci U S A. 2003;100:8532–7.

15. Wang Y, Liu L, Xia Z. Brain-derived neurotrophic factor stimulates the transcriptional and neuroprotective activity of myocyte-enhancer factor 2C through an ERK1/2-RSK2 signaling cascade. J Neurochem. 2007;102:957–66.

16. Lyons MR, Schwarz CM, West AE. Members of the myocyte enhancer factor 2 transcription factor family differentially regulate Bdnf transcription in response to neuronal depolarization. J Neurosci. 2012;32:12780–5.

17. Shieh PB, Hu SC, Bobb K, Timmusk T, Ghosh A. Identification of a signaling pathway involved in calcium regulation of BDNF expression. Neuron. 1998;20:727–40.

18. Tao X, Finkbeiner S, Arnold DB, Shaywitz AJ, Greenberg ME. Ca2+ influx regulates BDNF transcription by a CREB family transcription factor-dependent mechanism. Neuron. 1998;20:709–26.

19. Deogracias R, Espliguero G, Iglesias T, Rodriguez-Pena A. Expression of the neurotrophin receptor trkB is regulated by the cAMP/CREB pathway in neurons. Mol Cell Neurosci. 2004;26:470–80.

20. Lau GC, Saha S, Faris R, Russek SJ. Up-regulation of NMDAR1 subunit gene expression in cortical neurons via a PKA-dependent pathway. J Neurochem. 2004;88:564–75.

21. Biffo S, Offenhauser N, Carter BD, Barde YA. Selective binding and internalisation by truncated receptors restrict the availability of BDNF during development. Development. 1995;121:2461–70.

22. Alderson RF, Curtis R, Alterman AL, Lindsay RM, DiStefano PS. Truncated TrkB mediates the endocytosis and release of BDNF and neurotrophin-4/5 by rat astrocytes and schwann cells in vitro. Brain Res. 2000;871:210–22.

23. Rose CR, Blum R, Pichler B, Lepier A, Kafitz KW, Konnerth A. Truncated TrkB-T1 mediates neurotrophin-evoked calcium signalling in glia cells. Nature. 2003;426:74–8.

24. Aroeira RI, Sebastiao AM, Valente CA. BDNF, via truncated TrkB receptor, modulates GlyT1 and GlyT2 in astrocytes. Glia. 2015;63:2181–97.

25. Vaz SH, Jorgensen TN, Cristovao-Ferreira S, Duflot S, Ribeiro JA, Gether U, et al. Brain-derived neurotrophic factor (BDNF) enhances GABA transport by modulating the trafficking of GABA transporter-1 (GAT-1) from the plasma membrane of rat cortical astrocytes. J Biol Chem. 2011;286:40464–76.

26. Holt LM, Hernandez RD, Pacheco NL, Torres Ceja B, Hossain M, Olsen ML. Astrocyte morphogenesis is dependent on BDNF signaling via astrocytic TrkB.T1. Elife. 2019;8:e44667.

27. Ohira K, Kumanogoh H, Sahara Y, Homma KJ, Hirai H, Nakamura S, et al. A truncated tropomyosin-related kinase B receptor, T1, regulates glial cell morphology via Rho GDP dissociation inhibitor 1. J Neurosci. 2005;25:1343–53.

28. Ugalde-Triviño L, Tejeda GS, Esteban-Ortega GM, Díaz-Guerra M. A brain-accessible peptide modulates stroke inflammatory response and neurotoxicity by targeting BDNF-receptor TrkB-T1 specific interactome. Theranostics. 2025;15:4654–72.

29. Vidaurre OG, Gascón S, Deogracias R, Sobrado M, Cuadrado E, Montaner J, et al. Imbalance of neurotrophin receptor isoforms TrkB-FL/TrkB-T1 induces neuronal death in excitotoxicity. Cell Death Dis. 2012;3:e256.

30. Gomes JR, Costa JT, Melo CV, Felizzi F, Monteiro P, Pinto MJ, et al. Excitotoxicity downregulates TrkB.FL signaling and upregulates the neuroprotective truncated TrkB receptors in cultured hippocampal and striatal neurons. J Neurosci. 2012;32:4610–22.

31. Tejeda GS, Diaz-Guerra M. Integral characterization of defective BDNF/TrkB signalling in neurological and psychiatric disorders leads the way to new therapies. Int J Mol Sci. 2017;18:268.

32. Ferrer I, Krupinski J, Goutan E, Marti E, Ambrosio S, Arenas E. Brain-derived neurotrophic factor reduces cortical cell death by ischemia after middle cerebral artery occlusion in the rat. Acta Neuropathol. 2001;101:229–38.

33. Colombo E, Bacigaluppi M, Bartoccetti M, Triolo D, Bassani C, Bergamaschi A, et al. Astrocyte TrkB promotes brain injury and edema formation in ischemic stroke. Neurobiol Dis. 2024;201:106670.

34. Dorsey SG, Renn CL, Carim-Todd L, Barrick CA, Bambrick L, Krueger BK, et al. In vivo restoration of physiological levels of truncated TrkB.T1 receptor rescues neuronal cell death in a trisomic mouse model. Neuron. 2006;51:21–8.

35. Cao T, Matyas JJ, Renn CL, Faden AI, Dorsey SG, Wu J. Function and mechanisms of truncated BDNF receptor TrkB.T1 in neuropathic pain. Cells. 2020;9:1194.

36. Matyas JJ, O’Driscoll CM, Yu L, Coll-Miro M, Daugherty S, Renn CL, et al. Truncated TrkB.T1-mediated astrocyte dysfunction contributes to impaired motor function and neuropathic pain after spinal cord injury. J Neurosci. 2017;37:3956–71.

37. Choi DW. Glutamate neurotoxicity and diseases of the nervous system. Neuron. 1988;1:623–34.

38. Tejeda GS, Ayuso-Dolado S, Arbeteta R, Esteban-Ortega GM, Vidaurre OG, Diaz-Guerra M. Brain ischaemia induces shedding of a BDNF-scavenger ectodomain from TrkB receptors by excitotoxicity activation of metalloproteinases and gamma-secretases. J Pathol. 2016;238:627–40.

39. Tejeda GS, Esteban-Ortega GM, San Antonio E, Vidaurre ÓGG, Díaz-Guerra M. Prevention of excitotoxicity-induced processing of BDNF receptor TrkB-FL leads to stroke neuroprotection. EMBO Mol Med. 2019;11:e9950.

40. Esteban-Ortega GM, Torres-Campos E, Diaz-Guerra M. Retrograde transport of neurotrophin receptor TrkB-FL induced by excitotoxicity regulates Golgi stability and is a target for stroke neuroprotection. Cell Death Dis. 2025;16:659.

41. Lee YJ, Ch’ng TH. RIP at the Synapse and the Role of Intracellular Domains in Neurons. Neuromolecular Med. 2020;22:1–24.

42. Martin-de-Saavedra MD, Santos MD, Penzes P. Intercellular signaling by ectodomain shedding at the synapse. Trends Neurosci. 2022;45:483–98.

43. De Strooper B, Annaert W. Novel research horizons for presenilins and gamma-secretases in cell biology and disease. Annu Rev Cell Dev Biol. 2010;26:235–60.

44. Hallaq R, Volpicelli F, Cuchillo-Ibanez I, Hooper C, Mizuno K, Uwanogho D, et al. The Notch intracellular domain represses CRE-dependent transcription. Cell Signal. 2015;27:621–9.

45. Choi DW. Glutamate neurotoxicity in cortical cell culture is calcium dependent. Neurosci Lett. 1985;58:293–7.

46. Choi DW, Maulucci-Gedde M, Kriegstein AR. Glutamate neurotoxicity in cortical cell culture. J Neurosci. 1987;7:357–68.

47. Skowronska K, Obara-Michlewska M, Zielinska M, Albrecht J. NMDA receptors in astrocytes: in search for roles in neurotransmission and astrocytic homeostasis. Int J Mol Sci. 2019;20:309.

48. Milletti F. Cell-penetrating peptides: classes, origin, and current landscape. Drug Discov Today. 2012;17:850–60.

49. Regberg J, Eriksson JN, Langel U. Cell-penetrating peptides: from cell cultures to in vivo applications. Front Biosci (Elite Ed). 2013;5:509–16.

50. Ayuso-Dolado S, Esteban-Ortega GM, Vidaurre OG, Diaz-Guerra M. A novel cell-penetrating peptide targeting calpain-cleavage of PSD-95 induced by excitotoxicity improves neurological outcome after stroke. Theranostics. 2021;11:6746–65.

51. Lonze BE, Ginty DD. Function and regulation of CREB family transcription factors in the nervous system. Neuron. 2002;35:605–23.

52. Linseman DA, Bartley CM, Le SS, Laessig TA, Bouchard RJ, Meintzer MK, et al. Inactivation of the myocyte enhancer factor-2 repressor histone deacetylase-5 by endogenous Ca(2+) //calmodulin-dependent kinase II promotes depolarization-mediated cerebellar granule neuron survival. J Biol Chem. 2003;278:41472–81.

53. Woronicz JD, Lina A, Calnan BJ, Szychowski S, Cheng L, Winoto A. Regulation of the Nur77 orphan steroid receptor in activation-induced apoptosis. Mol Cell Biol. 1995;15:6364–76.

54. Zafra F, Castrén E, Thoenen H, Lindholm D. Interplay between glutamate and gamma-aminobutyric acid transmitter systems in the physiological regulation of brain-derived neurotrophic factor and nerve growth factor synthesis in hippocampal neurons. Proc Natl Acad Sci U S A. 1991;88:10037–41.

55. Burtrum D, Silverstein FS. Excitotoxic injury stimulates glial fibrillary acidic protein mRNA expression in perinatal rat brain. Exp Neurol. 1993;121:127–32.

56. Lin J, Yu Z, Gao X. Advanced noninvasive strategies for the brain delivery of therapeutic proteins and peptides. ACS Nano. 2024;18:22752–79.

57. Hong S, Piao J, Hu J, Liu X, Xu J, Mao H, et al. Advances in cell-penetrating peptide-based nose-to-brain drug delivery systems. Int J Pharm. 2025;678:125598.

58. Wu J, Roesger S, Jones N, Hu CJ, Li SD. Cell-penetrating peptides for transmucosal delivery of proteins. J Control Release. 2024;366:864–78.

59. Fumagalli S, Perego C, Ortolano F, De Simoni MG. CX3CR1 deficiency induces an early protective inflammatory environment in ischemic mice. Glia. 2013;61:827–42.

60. Stence N, Waite M, Dailey ME. Dynamics of microglial activation: a confocal time-lapse analysis in hippocampal slices. Glia. 2001;33:256–66.

61. Heindl S, Gesierich B, Benakis C, Llovera G, Duering M, Liesz A. Automated morphological analysis of microglia after stroke. Front Cell Neurosci. 2018;12:106.

62. Kikhia M, Schilling S, Herzog ML, Livne M, Semtner M, Tay TL, et al. Multicolor fate mapping of microglia reveals polyclonal proliferation, heterogeneity, and cell-cell interactions after ischemic stroke in mice. Nature Commun. 2025;16:8294.

63. Fonseca-Gomes J, Jerónimo-Santos A, Lesnikova A, Casarotto P, Castrén E, Sebastião AM, et al. TrkB-ICD fragment, originating from BDNF receptor cleavage, is translocated to cell nucleus and phosphorylates nuclear and axonal proteins. Front Mol Neurosci. 2019;12:4.

64. Gu Y, Zhou C, Piao Z, Yuan H, Jiang H, Wei H, et al. Cerebral edema after ischemic stroke: Pathophysiology and underlying mechanisms. Front Neurosci. 2022;16:988283.

65. Seillier C, Lesept F, Toutirais O, Potzeha F, Blanc M, Vivien D. Targeting NMDA receptors at the neurovascular unit: past and future treatments for central nervous system diseases. Int J Mol Sci. 2022;23:10336.

66. Arumugam TV, Chan SL, Jo DG, Yilmaz G, Tang SC, Cheng A, et al. Gamma secretase-mediated notch signaling worsens brain damage and functional outcome in ischemic stroke. Nat Med. 2006;12:621–623.

67. Polavarapu R, An J, Zhang C, Yepes M. Regulated intramembrane proteolysis of the low-density lipoprotein receptor-related protein mediates ischemic cell death. Am J Pathol. 2008;172:1355–62.

68. Shimada IS, Borders A, Aronshtam A, Spees JL. Proliferating reactive astrocytes are regulated by Notch-1 in the peri-infarct area after stroke. Stroke. 2011;42:3231–7.

69. Su T, Li Z, Yang Y, Dai Y, Li Y, Zhao H. In vitro 3D models of neuron-astrocyte interactions. Biochem Biophys Rep. 2026;45:102400.

70. Tam S, Wear D, Morrone CD, Yu WH. The complexity of extracellular vesicles: Bridging the gap between cellular communication and neuropathology. J Neurochem. 2024;168:2391–422.

71. Wan H, Cui Y, Zeng Y, Hu J, Li M, Xiao Z. Microglia-Astroglia-Neuron network following stroke: Novel insight into extracellular vesicles communication. Brain Res Bull. 2025;231:111537.

72. Varnamkhasti BS, Jafari S, Taghavi F, Alaei L, Izadi Z, Lotfabadi A, et al. Cell-penetrating peptides: as a promising theranostics strategy to circumvent the blood-brain barrier for CNS diseases. Curr Drug Deliv. 2020;17:375–86.

73. Pirhaghi M, Mamashli F, Moosavi-Movahedi F, Arghavani P, Amiri A, Davaeil B, et al. Cell-penetrating peptides: promising therapeutics and drug-delivery systems for neurodegenerative diseases. Mol Pharm. 2024;21:2097–117.

