## Supplementary Information for "The intracellular region of truncated neurotrophin receptor TrkB-T1 promotes stroke-related effects in glial reactivity and neurotoxicity"

### Supplementary Figures

**Additional file 1: Table S1.** Reagents and resources.

**Additional file 2: Fig. S1.** Subcellular fractionation of cultured astrocytes treated with APMA.

**Additional file 3: Fig. S2.** Excitotoxicity-induced RIP contributes to neuronal death of primary cultures of cortical neurons.

**Additional file 4: Fig. S3.** Microglia morphological analysis.

**Additional file 5: Fig. S4.** UMAP plot representing microglial cells from all mice and groups of treatment.

**Additional file 6:** Segmentation code used for analysis.

**Supplementary Table 1.**

| REAGENT or RESOURCE | SOURCE | IDENTIFIER |
|---|---|---|
| ***Antibodies*** | | |
| Mouse monoclonal anti-**β-Actin** | Sigma-Aldrich | Cat#A5441<br>RRID: AB_10920058 |
| Rabbit polyclonal anti-**C3d** | Dako | Cat#A0063<br>RRID: AB_578478 |
| Rabbit polyclonal anti-**Calnexin** | Enzo Life Science | Cat#ADI-SPA-860<br>RRID:AB_10616095 |
| Rabbit monoclonal anti-**CREB** | Cell Signaling | Cat#9197<br>RRID:AB_331277 |
| Rabbit monoclonal ani-**GFAP** | Millipore | Cat#MAB360<br>RRID:AB_11212597 |
| Mouse monoclonal anti-**GFAP** | Millipore | Cat#G6171<br>RRID:AB_1840893 |
| Rabbit polyclonal anti-**Iba1** | Wako | Cat#019-19741<br>RRID: AB_839504 |
| Mouse monoclonal anti-RBFOX3/**NeuN** | Novus | Cat#NBP1-92693SS;<br>RRID:AB_1103747 |
| Rabbit polyclonal anti-**TrkB-FL (C-ter region)** | Abcam | Cat#ab18987<br>RRID: AB_444716 |
| Rabbit polyclonal anti-**TrkB-T1** (isoform-specific C-ter) | Custom made | |
| Goat anti-mouse IgG Alexa Fluor 546 | Molecular Probes | Cat#A-11030;<br>RRID:AB_144695 |
| Goat anti-mouse IgG Alexa Fluor 647 | Molecular Probes | Cat#A-21236<br>RRID:AB_2535805 |
| Goat anti-rabbit IgG Alexa Fluor 488 | Molecular Probes | Cat# A11034<br>RRID: AB_2576217 |
| Goat anti-rabbit IgG Alexa Fluor 546 | Molecular Probes | Cat# A11035<br>RRID: AB_143051 |
| Donkey anti-rabbit IgG-heavy and light chain-HRP | Bethyl | Cat#A120-108P;<br>RRID:AB_10892625 |
| Donkey anti-mouse IgG-heavy and light chain-HRP | Bethyl | Cat#A90-137P;<br>RRID:AB_1211460 |
| ***Chemicals*** | | |
| **APMA** (used at 100 μM) | Calbiochem | Cat#164610<br>CAS: 6283-24-5 |
| **Ara C** (used at 10 μM) | Sigma-Aldrich | Cat#C1768;<br>CAS: 147-94-4 |
| **B27** serum free supplement | Thermo Fisher | Cat#17504044 |
| **Cy5-Streptavidin** | Palex Medical SA | Cat#SA-1500-1 |
| **DAPI** (used at 0.5 or 5 μg/ml) | Molecular Probes | Cat#D1306 |
| **EGF** (Human epidermal growth factor recombinant prot.**)** | Thermo Fisher | Cat# PHG0311 |
| b**FGF** (Human basic fibroblast growth factor recomb. prot.) | Thermo Fisher | Cat#13256-029 |
| **Fluorescein Avidin** D | Vector Laboratories | Cat#A 2001 |
| **Glycine** (used at 10 μM) | Bio-Rad | Cat#161-0718 |
| **Laminin** (used at 4 μg/ml) | Sigma-Aldrich | Cat#L2020;<br>CAS: 114956-81-9 |
| **MTT** (used at 0.5 mg/ml) | Sigma-Aldrich | Cat#M5655;<br>CAS: 298-93-1 |
| **NMDA** (used at 100 μM) | Tocris | Cat#0114;<br>CAS: 6384-92-5 |

| | | |
|---|---|---|
| **PhosSTOP** phosphatases inhibitor cocktail tablets | Roche | Cat#04 906 837 001 |
| **Poly-L-Lysine** (used at 100 µg/ml) | Sigma-Aldrich | Cat#P1524;<br>CAS: 25988-63-0 |
| **Prolong** Diamond antifade reagent | Molecular Probes | Cat#P36970 |
| Complete **protease inhibitor** cocktail tablets | Roche | Cat#11 697 498 001 |
| ***Peptides*** | | |
| **Bio-LTT1$_{Ct}$** (Biotin-*YGRKKRRQRRR*KLARHSKFGMKGFVLFHKIPLDG) | GenScript | N/A |
| **Bio-TMyc** (Biotin-*YGRKKRRQRRR*AEEQKLISEEDLLR) | GenScript | N/A |
| ***Primer sequences for real-time PCR*** | | |
| GluN1-F (all isoforms): TCCACCAAGAGCCCTTCGTG | Sigma-Aldrich | N/A |
| GluN1-R (all isoforms): GCCCCGTACAGATCACCTTC | Sigma-Aldrich | N/A |
| GluN2A-F: ACGACTGGGACTACAGCCTG | Sigma-Aldrich | N/A |
| GluN2A-R: CTTCTCTGCCTGCCCATAGC | Sigma-Aldrich | N/A |
| TrkB-FL-F (isoform specific): TATCTTCACCCACCTCAAAC | Sigma-Aldrich | N/A |
| TrkB-FL-R (isoform specific): GAGAGACTTGACCTGAGCAC | Sigma-Aldrich | N/A |
| TrkB-T1-F (isoform specific): GGGGCTGTGCTGCTTGGT | Sigma-Aldrich | N/A |
| TrkB-T1-R (isoform specific): GCTGCGGACATCTTTGGAGA | Sigma-Aldrich | N/A |
| BDNF-F (all isoforms): TGGCTGACACTTTTGAGCAC | Sigma-Aldrich | N/A |
| BDNF-R (all isoforms): GTTTGCGGCATCCAGGTAAT | Sigma-Aldrich | N/A |
| GAPDH-F: TTGCCATCAACGACCCCTTC | Sigma-Aldrich | N/A |
| GAPDH-R: GCCTTGACTGTGCCGTTGAA | Sigma-Aldrich | N/A |
| NSE-F: ACAGAATGGGGCTGTGTACC | Sigma-Aldrich | N/A |
| NSE-R: TGGCAACTGTGGGACATGGC | Sigma-Aldrich | N/A |
| ***Critical Commercial Assays*** | | |
| BCA Protein Assay Kit | Thermo Fisher | Cat# 23225 |
| Clarity Western ECL Blotting Substrate | BioRad | Cat# 1705060 |
| Lipofectamine 2000 | Life Technologies | Cat#11668019 |
| ***Experimental Models: Organisms/Strains*** | | |
| Balb/c inbred mice (Balb/cOlaHsd) | Janvier Labs SAS | N/A |
| Wistar Rat embryos (E18) | In site facility | N/A |
| ***Recombinant DNA*** | | |
| **pCRE** (25-mer oligonucleotide with the sequence of two TrkB CREs subcloned into pTK-Luc) | (Deogracias et al., 2004) | N/A |
| **pMEF2** (pRSRF; -307 to -242 of *Nur77* promoter with two MEF2 sites subcloned into pGL2-basic) | (Woronicz et al., 1995) | N/A |
| **pMEF2mut** (two inactivating point mutations in each MEF2 site of pMEF2) | (Woronicz et al., 1995) | N/A |
| ***Software*** | | |
| Adobe InDesign | | RRID:SCR_021799 |
| Adobe Photoshop | | RRID:SCR_014199 |
| GraphPad Prism | | RRID:SCR_002798 |
| ImageJ | imagej.net/ | RRID:SCR_003070 |
| R Project for Statistical Computing | www.r-project.org | RRID:SCR_001905 |
| AIVIA | version: 15.0.0.42239 | |
| ***Other*** | | |
| Tissue-Tek O.C.T Compound | Sakura | Cat#4583 |

| | | |
|---|---|---|
| Protran Western blotting nitrocellulose membrane | GE Healthcare | Cat#GE10600002; CAS: 9004-70-0 |

Key reagents and resources with indication of sources and identifiers.

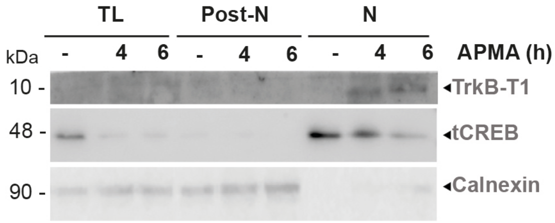
TL
Post-N
N
kDa
-
4
6
-
4
6
-
4
6
APMA (h)
10 -
48 -
90 -
TrkB-T1
tCREB
Calnexin

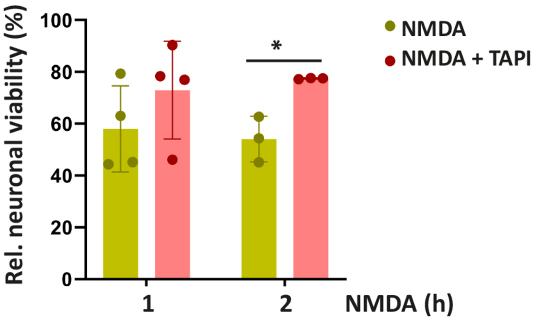

Rel. neuronal viability (%)
100
80
60
40
20
0
*
NMDA
NMDA + TAPI
1
2
NMDA (h)

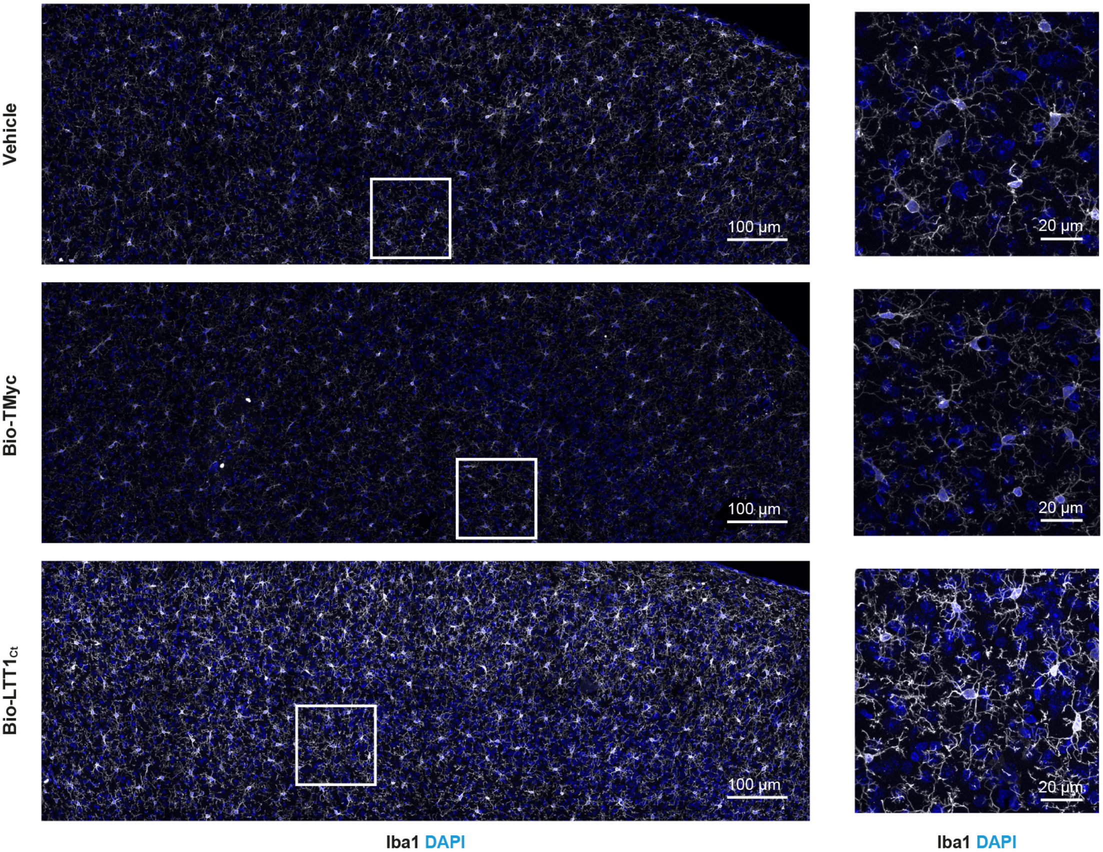
Vehicle
Bio-TMyc
Bio-LTT1Ct
100 µm
20 µm
100 µm
20 µm
100 µm
20 µm
Iba1 DAPI
Iba1 DAPI

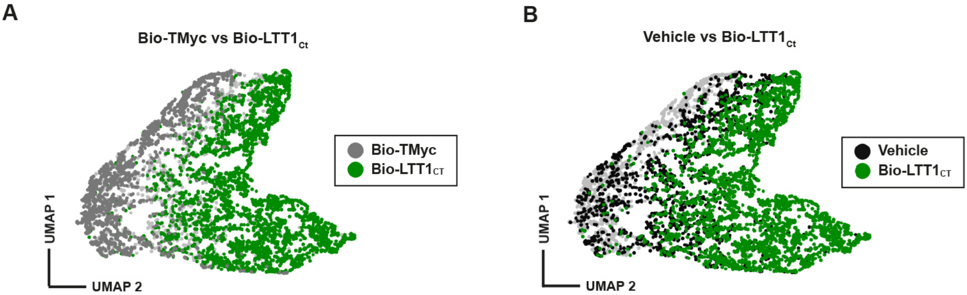
A
Bio-TMyc vs Bio-LTT1$_{Ct}$
Bio-TMyc
Bio-LTT1$_{CT}$
UMAP 1
UMAP 2
B
Vehicle vs Bio-LTT1$_{Ct}$
Vehicle
Bio-LTT1$_{CT}$
UMAP 1
UMAP 2

```
<?xml version="1.0" encoding="utf-8"?>
<RecipeApplyState>
  <SoftwareVersion>15.0.0.42239</SoftwareVersion>
  <SerializedVersion>1</SerializedVersion>
  <DeepLearningModelPath />
  <ChannelInputState>
    <PinID>6ff29774-9810-4fb7-abe2-e67642c5a517</PinID>
    <ChannelIndex>1</ChannelIndex>
  </ChannelInputState>
  <RecipeSettingsState>
    <RecipeName>3D Object Analysis - Meshes</RecipeName>
    <RecipeID>ddc70a23-5121-4110-bcc6-0599e3b1d67e</RecipeID>
    <RecipeParameterSetState>
      <RecipeName>3D Object Analysis - Meshes</RecipeName>
      <ParameterSetName>Detection</ParameterSetName>
      <ParameterSetID>7ab108c8-ac0f-4601-9dc0-108707305e37</
ParameterSetID>
    </RecipeParameterSetState>
    <RecipeParameterSetState>
      <RecipeName>3D Object Analysis - Meshes</RecipeName>
      <ParameterSetName>Partition</ParameterSetName>
      <ParameterSetID>4f471a17-2942-4d0f-b919-0edf74e11c5d</
ParameterSetID>
      <RecipeParameterValue>
        <ParameterName>Max Object Radius</ParameterName>
        <ParameterID>27e90136-bdb0-4c51-b37f-c6b75ec5a57a</
ParameterID>
        <ParameterValue>200</ParameterValue>
      </RecipeParameterValue>
      <RecipeParameterValue>
        <ParameterName>Min Object Radius</ParameterName>
        <ParameterID>6883d75d-bdb7-459d-8213-3e6a1b8c1e65</
ParameterID>
        <ParameterValue>5</ParameterValue>
      </RecipeParameterValue>
      <RecipeParameterValue>
        <ParameterName>Mesh Smoothing Factor</ParameterName>
        <ParameterID>ffd11422-6b54-4837-93a9-e9e454358404</
ParameterID>
        <ParameterValue>0</ParameterValue>
      </RecipeParameterValue>
    </RecipeParameterSetState>
    <RecipeParameterSetState>
      <RecipeName>Skip Smooth Image</RecipeName>
      <ParameterSetName>Enhance Cells</ParameterSetName>
      <ParameterSetID>8e49dc27-1f25-4f3a-aa5c-9ac7384baf78</
ParameterSetID>
      <RecipeParameterValue>
        <ParameterName>Image Smoothing Filter Size</ParameterName>
        <ParameterID>baf5f5e3-23df-43f1-9fc0-82c087818e4b</
ParameterID>
        <ParameterValue>0</ParameterValue>
      </RecipeParameterValue>
    </RecipeParameterSetState>
```

```
    <RecipeParameterSetState>
      <RecipeName>Morphological Smoothing</RecipeName>
      <ParameterSetName>Smooth Cells</ParameterSetName>
      <ParameterSetID>52a8b407-b8d2-4cfe-9683-b24948746904</
ParameterSetID>
      <RecipeParameterValue>
        <ParameterName>Image Smoothing Filter Size</ParameterName>
        <ParameterID>8bc07246-a5c6-4f43-9f10-1fd0ac167bb3</
ParameterID>
        <ParameterValue>1</ParameterValue>
      </RecipeParameterValue>
    </RecipeParameterSetState>
    <RecipeParameterSetState>
      <RecipeName>Average Filter Smoothing</RecipeName>
      <ParameterSetName>Smooth Cells</ParameterSetName>
      <ParameterSetID>52a8b407-b8d2-4cfe-9683-b24948746904</
ParameterSetID>
      <RecipeParameterValue>
        <ParameterName>Image Smoothing Filter Size</ParameterName>
        <ParameterID>8bc07246-a5c6-4f43-9f10-1fd0ac167bb3</
ParameterID>
        <ParameterValue>9</ParameterValue>
      </RecipeParameterValue>
    </RecipeParameterSetState>
    <RecipeParameterSetState>
      <RecipeName>Gaussian Filter Smoothing</RecipeName>
      <ParameterSetName>Smooth Cells</ParameterSetName>
      <ParameterSetID>52a8b407-b8d2-4cfe-9683-b24948746904</
ParameterSetID>
      <RecipeParameterValue>
        <ParameterName>Image Smoothing Filter Size</ParameterName>
        <ParameterID>8bc07246-a5c6-4f43-9f10-1fd0ac167bb3</
ParameterID>
        <ParameterValue>9</ParameterValue>
      </RecipeParameterValue>
    </RecipeParameterSetState>
    <RecipeParameterSetState>
      <RecipeName>Median Filter Smoothing</RecipeName>
      <ParameterSetName>Smooth Cells</ParameterSetName>
      <ParameterSetID>52a8b407-b8d2-4cfe-9683-b24948746904</
ParameterSetID>
      <RecipeParameterValue>
        <ParameterName>Image Smoothing Filter Size</ParameterName>
        <ParameterID>8bc07246-a5c6-4f43-9f10-1fd0ac167bb3</
ParameterID>
        <ParameterValue>9</ParameterValue>
      </RecipeParameterValue>
    </RecipeParameterSetState>
    <RecipeParameterSetState>
      <RecipeName>Skip Remove Background</RecipeName>
      <ParameterSetName>Object Detection</ParameterSetName>
      <ParameterSetID>a5d14608-cc7d-436c-97c3-cc511ba2a08d</
ParameterSetID>
      <RecipeParameterValue>
```

```
        <ParameterName>Min Edge Intensity</ParameterName>
        <ParameterID>36d6211a-0944-4bc3-9ea6-e9e2f18236ae</
ParameterID>
        <ParameterValue>0.00784313725490196</ParameterValue>
      </RecipeParameterValue>
      <RecipeParameterValue>
        <ParameterName>Fill Holes Size</ParameterName>
        <ParameterID>333ec50c-2d60-4ee4-912d-f5864b73c9a3</
ParameterID>
        <ParameterValue>0</ParameterValue>
      </RecipeParameterValue>
    </RecipeParameterSetState>
    <RecipeParameterSetState>
      <RecipeName>Remove Background</RecipeName>
      <ParameterSetName>Object Detection</ParameterSetName>
      <ParameterSetID>a5d14608-cc7d-436c-97c3-cc511ba2a08d</
ParameterSetID>
      <RecipeParameterValue>
        <ParameterName>Average Object Radius</ParameterName>
        <ParameterID>ea132080-a499-40a1-9035-93f7ce144cdf</
ParameterID>
        <ParameterValue>20</ParameterValue>
      </RecipeParameterValue>
      <RecipeParameterValue>
        <ParameterName>Min Edge Intensity</ParameterName>
        <ParameterID>f1c400e1-d376-4fb4-ad27-27608242db0d</
ParameterID>
        <ParameterValue>0.12</ParameterValue>
      </RecipeParameterValue>
      <RecipeParameterValue>
        <ParameterName>Fill Holes Size</ParameterName>
        <ParameterID>1fea40e0-5167-4a03-b6ca-fd12703b5d4e</
ParameterID>
        <ParameterValue>0</ParameterValue>
      </RecipeParameterValue>
    </RecipeParameterSetState>
    <RecipeParameterSetState>
      <RecipeName>Apply Partition</RecipeName>
      <ParameterSetName>Partition</ParameterSetName>
      <ParameterSetID>aa76e840-fe29-4414-93c3-ecd47d3b9533</
ParameterSetID>
      <RecipeParameterValue>
        <ParameterName>Min Edge to Center Distance</ParameterName>
        <ParameterID>a62966fc-39de-43fa-ab97-7362df00cb07</
ParameterID>
        <ParameterValue>20</ParameterValue>
      </RecipeParameterValue>
    </RecipeParameterSetState>
    <RecipeParameterSetState>
      <RecipeName>Skip Partition</RecipeName>
      <ParameterSetName>Partition</ParameterSetName>
      <ParameterSetID>a949756a-c62e-4b2d-8c12-33620011ac7d</
ParameterSetID>
      <RecipeParameterValue>
```

```
        <ParameterName>Min Edge to Center Distance</ParameterName>
        <ParameterID>f7b6e05a-d0ad-441e-9bd6-7673bf246de5</
ParameterID>
        <ParameterValue>20</ParameterValue>
      </RecipeParameterValue>
    </RecipeParameterSetState>
    <SwitchableRecipeState>
      <RecipeName>Smoothing_WholeCell_switchable</RecipeName>
      <SelectedRecipeName>Morphological Smoothing</
SelectedRecipeName>
    </SwitchableRecipeState>
    <SwitchableRecipeState>
      <RecipeName>ImageEnhancementOptions</RecipeName>
      <SelectedRecipeName>Skip Remove Background</
SelectedRecipeName>
    </SwitchableRecipeState>
    <SwitchableRecipeState>
      <RecipeName>ObjectPartitionOptions</RecipeName>
      <SelectedRecipeName>Apply Partition</SelectedRecipeName>
    </SwitchableRecipeState>
    <RecipeOutputGroupState>
      <OutputGroupName>Cross Sections</OutputGroupName>
      <OutputGroupID>454067e5-9dce-4387-80eb-fb1ad7be2e01</
OutputGroupID>
      <IsEnabled>true</IsEnabled>
    </RecipeOutputGroupState>
  </RecipeSettingsState>
</RecipeApplyState>
```
